# OrthoGather: a local platform for orthology-based proteome comparisons and Gene Ontology enrichment

**DOI:** 10.64898/2026.01.30.702851

**Authors:** Carlos Vivas-Rodríguez, David Gomez Matallanas, Colm J. Ryan, Siobhán McClean, Olivier Dennler, Joanna Drabinska

## Abstract

Comparative proteomic analysis may reveal common and unique pathways regulated by the same stimulus across species using data from differential protein expression studies or curated protein sets. Functional annotations are key but vary in quality, as many proteins, particularly in prokaryotes and non-model eukaryotes, are poorly or inconsistently annotated, complicating comparative studies. Orthology inference provides a robust framework to address this, but currently available orthology and comparative genomics tools technical expertise, command-line use, and manual processing of complex outputs, creating barriers for researchers without computational training. We developed *OrthoGather*, a locally hosted web application that streamlines comparative proteomic analysis by integrating orthologous protein groups across species and Gene Ontology (GO) enrichment. It can integrate functional annotations from proteins within orthogroups to support the functional interpretation of protein sets, even when individual species lack comprehensive annotation. Its flexible design supports cross-species exploration of conserved and unique orthogroups across proteomes or user-defined protein sets, revealing functional patterns through orthogroup relationships. *OrthoGather* generates publication-ready and interactive outputs, including downloadable plots and data files, lowering barriers for researchers without computational expertise.

## Introduction

The incomplete functional annotation of proteins remains a major challenge in proteomics, particularly when interpreting biological significance of protein sets derived from experimental studies. This is especially challenging in prokaryotes or non-model eukaryotes, where annotations are commonly incomplete or inconsistent [1-5]. This variability complicates cross-species comparisons, limiting the identification of functionally equivalent proteins and shared or species-specific responses to stimuli [6-8]. Orthology inference addresses this limitation by identifying evolutionarily related proteins across species. Tools such as OrthoFinder enable distinguishing conserved and species-specific gene sets by clustering evolutionarily related proteins into orthogroups [9,10,13].

Numerous tools support orthology inference, comparative genomics, functional annotation, and visualisation (Supplementary Table S1), but most address individual workflow stages. Consequently, comparative analyses often require multiple tools. Additionally, many workflows rely on command-line execution, manual data preparation, or predefined reference datasets, producing complex outputs that can limit accessibility and complicate comparative analysis of experimentally derived protein sets, particularly for researchers without computational expertise.

To address these issues, we developed *OrthoGather*, a locally hosted web application streamlining comparative proteomic analysis. It retrieves reference proteomes from the UniProt database [6], executes OrthoFinder locally, and generates browser-based, downloadable figures and summary statistics. Gene Ontology (GO) enrichment supports functional interpretation of user-defined protein sets in a comparative, cross-species context [11,12]. GO annotations associated with proteins within orthogroups can provide functional context, even in poorly annotated species. OrthoGather can map experimentally derived protein sets (e.g. proteins changed in abundance following exposure to stress) to orthogroups across other species and use GO enrichment to explore the functions represented within those orthogroups.

### *OrthoGather* Development and Implementation

*OrthoGather* is developed in Python 3.11, using Flask as the backend framework, with an HTML and JavaScript frontend. Interaction is supported across modern web browsers without additional installation.

The application has a modular architecture with four components (Fig. 1). (1) *Proteome selection* - a predictive UniProt proteome search [6], using a local index of over one million UniProt proteomes searchable by scientific or common name, taxonomic synonym, or UniProt proteome identifier, followed by streamed FASTA download from the UniProt REST API with automatic retry on failure. (2) *Orthogroup inference* - OrthoFinder v2.5.5 [9,10] is executed internally via Python’s subprocess module using the -og flag, stopping after orthogroup clustering and omitting gene-tree and pairwise orthologue inference; only the Orthogroups directory and summary statistics are retained and reported to minimise storage (Supplementary File S1, Fig. S3), while empty or low-coverage proteomes are flagged. (3) *Orthogroup filtering and exploration* – filtering by selected species and user-defined UniProt identifiers, with interactive visualisation via Plotly.js [39] and UpSet plots [14], with export in CSV, XLSX, TSV, and JSON. (4) *Functional analysis using Gene Ontology* - Gene Ontology Annotation (GOA) files are retrieved from the European Bioinformatics Institute [15]. The foreground comprises a list of UniProt identifiers. The background can be defined as a custom UniProt identifier list or all proteins from species with available GOA files. The *Expand to full orthogroups* option expands each set to all proteins in orthogroups containing at least one selected entry. Enrichment analysis is performed using GOATOOLS [16] against the Gene Ontology [11,12]. *OrthoGather* runs as a locally hosted web application compatible with macOS, Linux and Windows (via Windows Subsystem for Linux). Guided installers automate dependency setup, eliminating command-line use. Internet access is required for proteome, GO annotation and optional catalogue retrieval; all computation is performed locally. Benchmarking on an Apple M4 (10 cores, 24 GB RAM) showed the example analysis completed in >3 min. Peak memory was ∼1 GB and the analysis added a measured ∼6 MB to the on-disk footprint. A multi-core machine with ≥8 GB RAM and ∼1 GB free disk is sufficient for comparable analysis.

**Figure 1.**
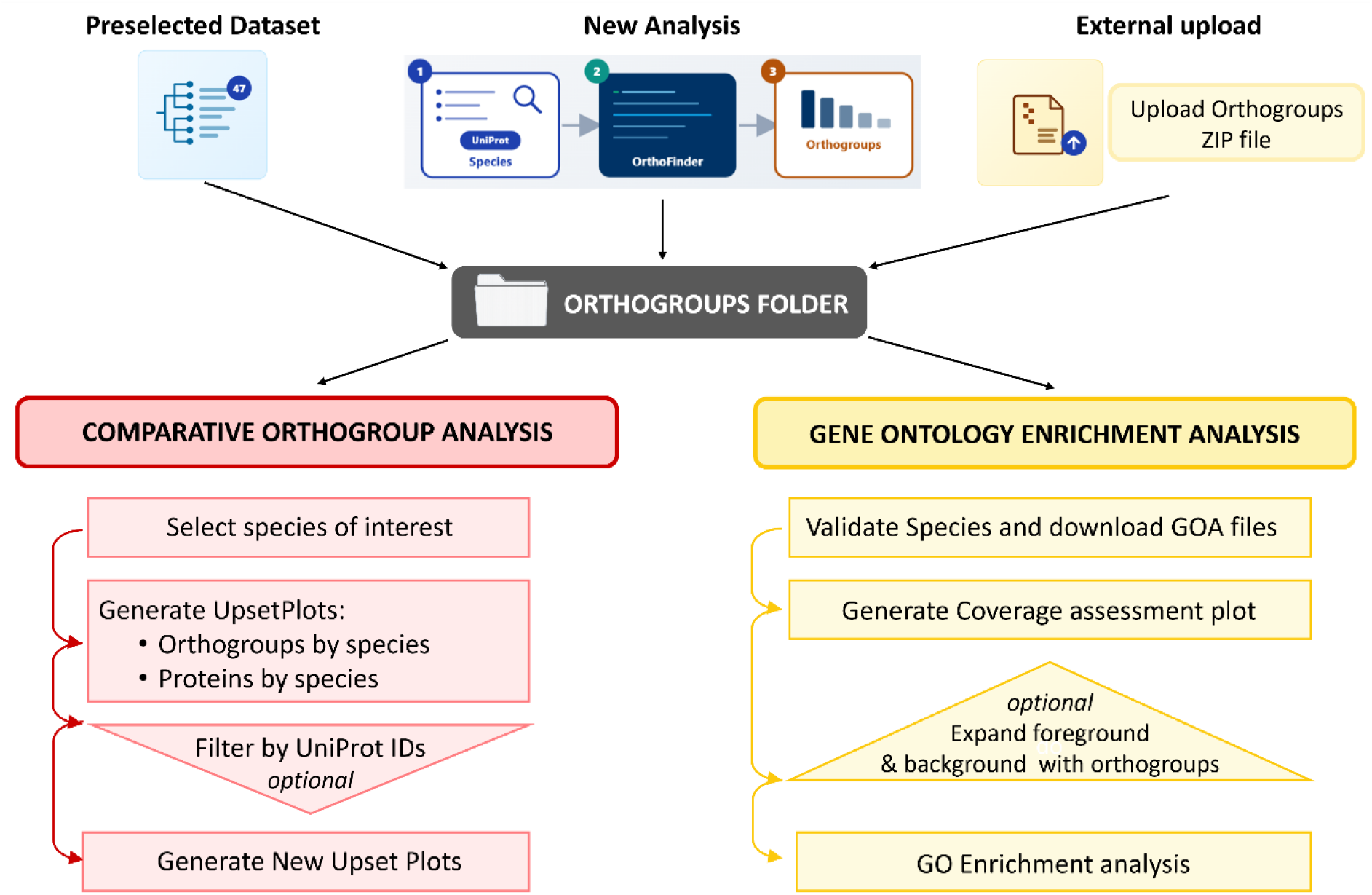
Overview of the *OrthoGather* workflow. All input modes: *External Data Upload, New Analysis*, and *Preselected Dataset* converge in the *Orthogroups* folder. The data can be explored through two modules: *Comparative Orthogroup Analysis* and *Gene Ontology Enrichment*

### *OrthoGather* Workflow and Output Overview

*OrthoGather* integrates comparative proteome analysis with GO enrichment, streamlining data acquisition and interpretation (Fig. 1). An example application, using a differential proteomics dataset from a study examining the response of *Mycolicibacterium smegmatis* (Msm) to sub-lethal concentrations of rifampicin [17], is provided in Supplementary File S1 and used throughout.

*OrthoGather* supports three input modes (Fig. 1; Supplementary File S1, Fig. S1). (1) *New Analysis*, an autocomplete search supporting scientific and common names, taxonomic synonyms, and UniProt proteome IDs, across over one million UniProt proteomes (Supplementary File S1, Fig. S2). A taxonomy-based tree built from NCBI Taxonomy lineages [18] visualises the classification of selected species and can be exported in Newick format (Fig. 2A). Selected reference and non-reference proteomes are automatically downloaded in FASTA format, and *OrthoFinder* infers orthogroups for subsequent analyses. (2) *Preselected Dataset*, 47 proteomes from 35 species across 19 genera associated with cystic fibrosis lung infections and/or antimicrobial resistance (Supplementary File S2), enabling familiarisation with the platform without proteome downloads or external computations. (3) *External Data Upload* accepts compressed *OrthoFinder* results containing the *Orthogroups* directory. OrthoXML relationships can be imported using OrthoXML-Tools [19], enabling reuse of external analyses and interoperability with existing orthology databases. Regardless of input mode, *OrthoGather* centralises processing in the *Orthogroups* folder, with temporary and auxiliary OrthoFinder files removed to optimise disk usage. Subsequently, users can perform *Comparative Orthogroup Analysis* and/or *Gene Ontology Enrichment Analysis*.

**Figure 2.**
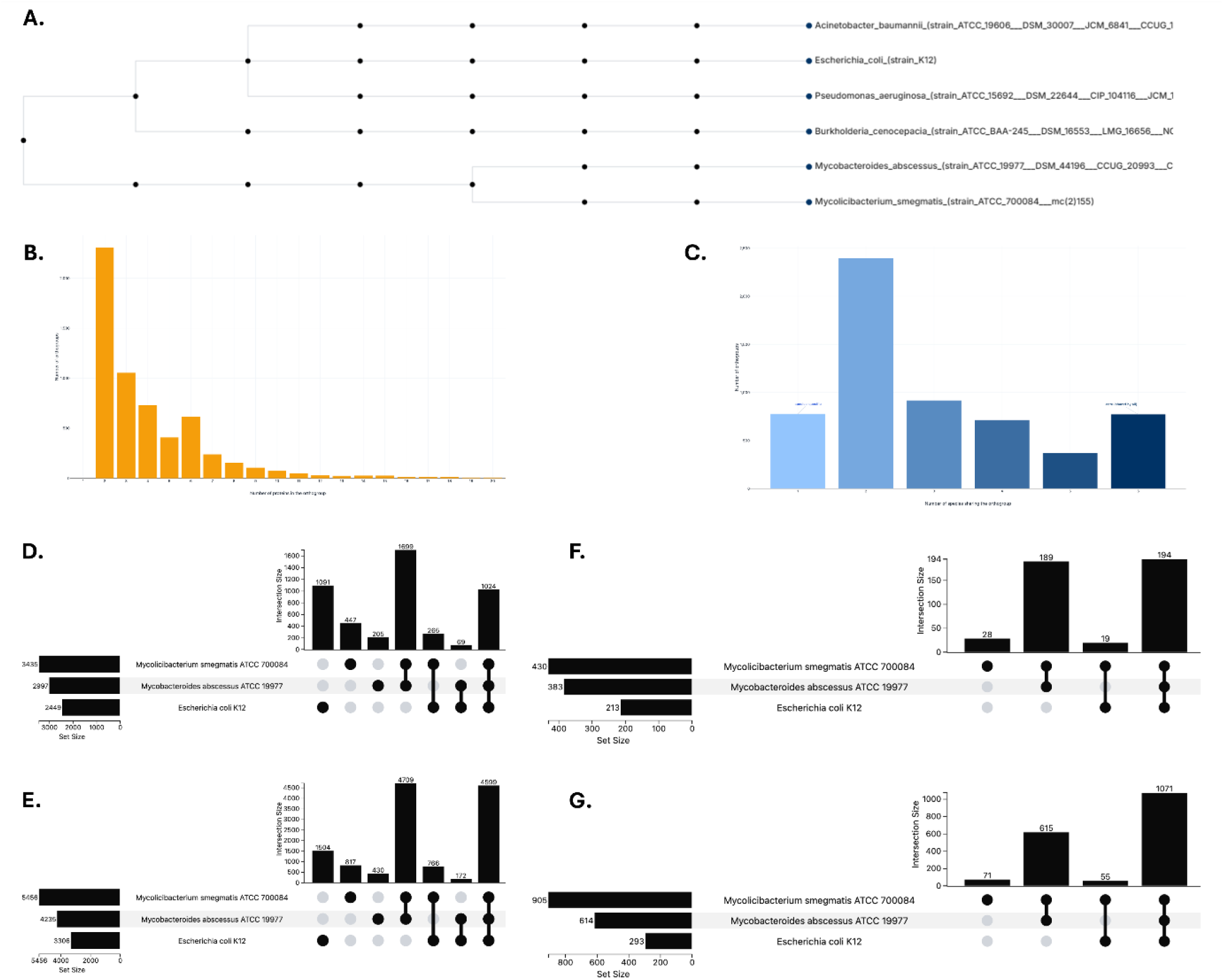
Comparative Orthogroup Analysis outputs. **(A)** Taxonomic dendrogram of the selected reference proteomes, generated on demand by *OrthoGather* from NCBI Taxonomy lineages [18] and exportable in Newick format. It provides a taxonomic overview to support species selection and represents the NCBI taxonomic classification of the species. *Mycolicibacterium smegmatis, Mycobacteroides abscessus, Acinetobacter baumannii, Burkholderia cenocepacia, Escherichia coli*, and *Pseudomonas aeruginosa* were selected. **(B)** Distribution of orthogroup sizes, showing the number of proteins within each orthogroup. **(C)** Distribution of orthogroup conservation across analysed species, indicating the number of species represented within each orthogroup. **(D–E)** Representative UpSet plots generated following species filtering of the example dataset, illustrating orthogroup and protein overlap among the selected species. The analysis was restricted to *M. smegmatis, M. abscessus* and *E. coli*. **(D)** UpSet plot showing the number of orthogroups in each species and their intersections. **(E)** UpSet plot showing the number of proteins within the orthogroup intersections. **(F–G)** Representative UpSet plots generated following filtering of the example dataset using the foreground protein set (Supplementary File S1, Table S5), retaining only orthogroups containing at least one selected protein. **(F)** UpSet plot showing the number of retained orthogroups in each species and their intersections. **(G)** UpSet plot showing the number of proteins within the retained orthogroup intersections. In all UpSet plots, the left bar chart shows the total number of orthogroups, or proteins associated with each species, while the upper bar chart shows the size of each exclusive intersection defined by the species combinations indicated below.

The *Comparative Orthogroup Analysis* module explores orthogroup composition across species (Fig. 2B–C) using two filtering phases. In the *Species filtering phase*, at least two species are selected (Supplementary File S1, Fig. S4), retaining only orthogroups containing at least one protein from the selected species, excluding those formed exclusively by non-selected taxa. Two UpSet plots show orthogroup distribution across species combinations (Fig. 2D) and the number of proteins in each intersection (Fig. 2E); hovering reveals the proportional contribution of each species. To illustrate the workflow, the *New Analysis* input mode was used with proteomes of *Mycolicibacterium smegmatis, Mycobacteroides abscessus, Acinetobacter baumannii, Burkholderia cenocepacia, Escherichia coli* and *Pseudomonas aeruginosa* (Supplementary File S1, Fig. S2), then limited to Mab, Msm and *E. coli* for UpSet plot generation. The *Protein filtering phase* further narrows the analysis using a custom UniProt ID list (Supplementary File S1, Fig. S5). *OrthoGather* retains orthogroups with at least one provided ID, generating new UpSet plots (Fig. 2F-G). Unmatched identifiers are classified as unassigned genes, present in the analysed proteomes but not assigned to an orthogroup by OrthoFinder, or absent from the analysis. Of the 491 proteins differentially expressed in Msm following rifampicin treatment (Supplementary File S1), 467 were mapped to 430 orthogroups, while 24 were unassigned. Retained orthogroups and proteins were visualised to identify shared and species-specific orthogroups (Supplementary File S1, Fig. S5B–C). Plot data can be exported in CSV, XLSX, TSV, and JSON and include UniProt identifiers of proteins for each orthogroup and species, enabling identification of homologous proteins, even when names differ.

The *Gene Ontology Enrichment Analysis* module enables functional interpretation of filtered and unfiltered orthogroups using annotations from species represented in the GOA repository [15]. The workflow has three phases. First, in the *Species validation phase, OrthoGather* checks analysed species for corresponding GOA files using a multi-step algorithm expanding common names and abbreviated binomials and applying three-tier matching against the UniProt proteome catalogue. Only species with GOA annotations contribute to downstream GO analysis, while species without annotations remain in the orthogroup structure but contribute no GO information. Manual entry of a taxon ID or species name is possible. In the analysed example, all selected species were automatically matched to GOA files (Supplementary File S1, Fig. S6A). Second, in the *Functional coverage phase, OrthoGather* quantifies the proportion of proteins in each orthogroup associated with at least one GO term. This provides downloadable GO annotation coverage data. Coverage is displayed as a stacked histogram (Fig. 3A), allowing overall annotation level assessment. Low coverage suggests that additional well-annotated species could be included. In the analysed example, GOA files were available for all six species. Of the 5,949 orthogroups, 4,638 (78.0%) contained at least one annotated protein, while the rest contained none and were not considered in the functional enrichment analysis. Among the annotated orthogroups, coverage had a median of 100% and a mean of 93.8% (Fig. 3A–B), Supplementary File S1, Fig. S6B–C).

**Figure 3.**
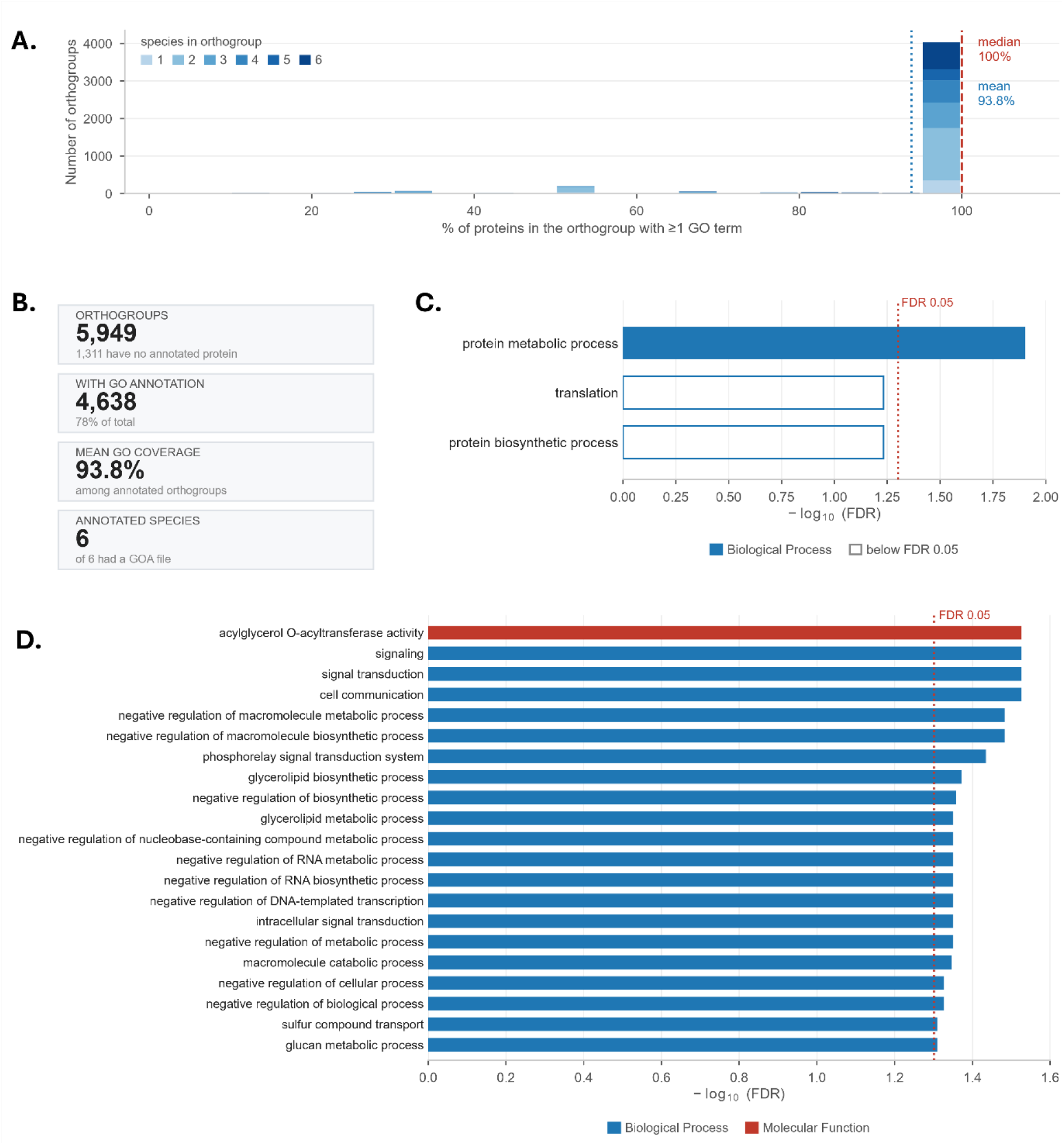
Gene Ontology Enrichment Analysis outputs. **(A)** Distribution of GO annotation coverage across orthogroups, showing the proportion of proteins associated with at least one GO term, stacked by the number of species. Vertical lines mark the median (100%) and the mean (93.8%) coverage among annotated orthogroups. **(B)** Summary of GO annotation coverage across the analysed orthogroups. **(C–D)** GO enrichment results for the example dataset. Proteins increased in abundance (n = 185; 162 tested) and proteins decreased in abundance (n = 306; 250 tested) were analysed as separate foregrounds against a common background of 3,181 proteins, of which 2,470 were GO-annotated. **(C)** GO enrichment results for proteins increased in abundance. **(D)** GO enrichment results for proteins decreased in abundance. An FDR threshold of 0.05 and a minimum GO depth of 2 were applied to both analyses.

Lastly, in the *Enrichment phase*, two protein sets are defined: *foreground*, containing proteins of interest, and background, against which the enrichment is evaluated. The background can be defined as all proteins from species with available GOA files or a custom UniProt ID list. The *Expand to full orthogroups* option expands each set to all proteins in orthogroups containing at least one selected entry. The example dataset was analysed without expansion, so the enrichment describes the Msm proteins reported by the original study. As analysis of the 491 differentially abundant proteins from the analysed example dataset returned only one significant term (protein maturation), they were split into two foregrounds, 185 increased and 306 decreased in abundance, against a common background; separate analysis, with the false discovery rate (FDR) controlled within each, identified 21 significant terms for the decreased-in-abundance proteins and one for the increased-in-abundance proteins. The *background* represents the complete protein set detected in the experiment, here, 3,181 Msm proteins detected by mass spectrometry (Supplementary File S1, Fig. S6D-F). The analysis uses GOATOOLS v1.6.4 [16] and go-basic.obo release 2026-07-26; each term is tested with a one-sided Fisher’s exact test (SciPy [40]) for over-representation in the foreground relative to the background. Annotations are propagated through the ontology *via* is_a and part_of relations (True Path Rule). The population comprises background proteins carrying at least one GO annotation, with the same annotation requirement applied to the foreground. As in PANTHER’s overrepresentation test [29], a single population is used for Biological Process, Cellular Component and Molecular Function. Terms below the selected minimum GO depth, and terms annotated to fewer than 10 or more than 500 proteins of the annotated background (the gene-set size limits used in clusterProfiler and WebGestalt [37,38]), are excluded before correction. The Benjamini–Hochberg FDR [20] is then computed across the retained terms, pooled across the three GO namespaces. Significant results are displayed as bar plots (Fig. 3C–D). Adjustable parameters include p-value threshold, ontology depth, maximum number of displayed terms, evidence-code stringency (all, non-IEA, or experimental only), inclusion of all or only single-copy orthogroups and counting mode (per protein or per orthogroup), enabling users to tailor the analysis and outputs to their needs. Results are presented in an interactive table exportable in CSV, TSV and JSON, alongside a multi-sheet XLSX workbook with full analysis results.

In the analysed example, an FDR threshold of 0.05 and a minimum GO depth of 2 were applied. The increased-in-abundance protein set returned one significant term (Fig. 3C). The decreased-in-abundance protein set returned 21 significant terms (Fig. 3D). 98.3% of the GO annotations available for Msm are electronic (IEA), reflecting predicted functional annotations. The resulting enrichment analysis highlighted biological processes associated with the response to rifampicin (Supplementary File S1, Fig. S7–S8).

## Discussion

A wide range of tools supports orthology inference, comparative genomics, and functional annotation, with most focusing on specific workflow stages (Supplementary Table S1). Tools such as OrthoFinder, OrthoMCL, Proteinortho, InParanoid, SonicParanoid, and Broccoli focus on orthology inference from user-provided proteomes [10,21-25], whereas OMA, OrthoDB, Ensembl Compara, PANTHER and MBGD provide curated orthology databases supporting exploration of precomputed orthologous groups, gene families, and genomic relationships for predefined species [26-30]. Functional annotation and enrichment are addressed by eggNOG, STRING, KEGG Orthology, ShinyGO and Blast2GO [5,31-34]. Consequently, comparative analysis of experimentally derived protein sets requires multiple resources, manual preparation of intermediate files and computational expertise. *OrthoGather* reduces technical barriers while remaining compatible with established orthology workflows, enabling interoperability with external orthology databases [9,10,19].

OrthoVenn3 supports some functionality similar to *OrthoGather*, including orthology analysis and comparative visualisation through a web interface [35]. However, it focuses on orthogroup identification and comparison, while *OrthoGather* combines automated proteome retrieval, *de novo* orthology inference, analysis of user-defined protein sets and GO enrichment within a single application. While OrthoVenn3 provides a built-in database of 733 species or accepts uploaded FASTA files [35], *OrthoGather* provides access to over one million UniProt proteomes together with per-orthogroup counting and evidence-code filtering. *OrthoGather* does not include collinearity analysis, gene family expansion and contraction analysis or maximum-likelihood species tree inference [35].

*OrthoGather* does not provide additional orthology information but makes comparative orthology-based analyses more accessible to researchers without computational experience by integrating analytical steps into a single reproducible workflow. Data provenance tracking, an integrated tutorial, downloadable datasets, and publication-ready figures reduce manual processing and support reproducibility, while analysis history allows users to save, browse, reproduce, and compare runs. Local hosting keeps all data on the user’s machine, avoiding privacy concerns associated with uploading unpublished datasets to external servers and enabling offline use once installed.

Although developed primarily for comparative proteomics, *OrthoGather* can support transcriptomic and genomic datasets, provided proteins or genes can be associated with reference proteomes for orthology inference. Applications include comparing differentially expressed proteins or genes across species, identification of conserved and species-specific responses to environmental or experimental conditions, analysis of orthologues of individual proteins or gene families, and interpretation of whole-genome sequencing or other high-throughput datasets. By integrating orthology relationships with comparative and functional analyses, *OrthoGather* supports investigation of evolutionary conservation and generation of biologically meaningful hypotheses across diverse organisms.

Several factors need to be considered when interpreting the outputs of OrthoGather. The comparative analyses depend on orthogroup inference generated by OrthoFinder and the completeness of the selected reference proteomes. Furthermore, OrthoGather does not implement alternative orthology inference algorithms, hierarchical orthologous group models, or detailed evolutionary analyses available in specialised resources [21-30]. This orthogroup framework also provides the basis for downstream functional analysis, in which OrthoGather uses orthogroups as an evolutionary framework for comparison while GO enrichment uses existing GO annotations associated with proteins included in the analysis. When complete orthogroups are included, annotations from orthologous proteins contribute to enrichment, increasing the representation of existing GO terms and providing additional functional context, particularly for poorly annotated species. Thus, the analysis can identify GO terms that are overrepresented among evolutionarily related proteins within the selected orthogroups and proteomes. Importantly, this does not constitute orthology-based functional annotation or inference of protein function from orthogroup membership; annotations are not transferred or assigned to unannotated proteins, and OrthoGather does not implement phylogeny-based annotation propagation such as PAINT [36]. GO enrichment also depends on the availability and quality of Gene Ontology Annotation (GOA) datasets, and species lacking GOA annotations cannot contribute functional information. Interpretation also depends on the evidence behind the underlying annotations. Finally, OrthoGather is intended for focused, user-defined comparisons rather than large-scale orthology analyses. Computational performance scales with the number and size of the selected proteomes and the available user hardware, with the main computational bottleneck arising from OrthoFinder inference.

To conclude, *OrthoGather* is a locally hosted web application that integrates orthology inference, comparative orthogroup analysis and GO enrichment into a single browser-based workflow. Rather than replacing specialised orthology and annotation resources, *OrthoGather* complements them by lowering technical barriers to comparative proteome analysis for researchers without extensive computational expertise, particularly for prokaryotes and other poorly annotated non-model organisms. Applied to a published differential proteomics dataset, *OrthoGather* recovered the functional changes reported by the original study, including reduced glycerolipid biosynthesis, two-component signalling and transcriptional regulation, through GO enrichment, while orthogroup analysis identified conserved and species-specific patterns across bacterial species with varying levels of functional annotation and without requiring command-line tools or manual data processing.

## Supporting information

Supplementary File S1 - Example dataset analysis

Supplementary File S2 - Preselected dataset

Supplementary Table S1 - Tool comparison

## Abbreviations

API: application programming interface
CSV: comma-separated values
FASTA: Fast-All sequence format
GOA: Gene Ontology Annotation
ID: identifier
IEA: inferred from electronic annotation
JSON: JavaScript Object Notation
Mab: *Mycobacteroides abscessus*
Msm: *Mycolicibacterium smegmatis*
NCBI: National Centre for Biotechnology Information
RAM: random-access memory
REST: Representational State Transfer
TSV: tab-separated values
XLSX: Microsoft Excel Open XML (Extensible Markup Language) spreadsheet format

## Author Contributions

Joanna Drabinska conceived and initiated the project, secured funding, provided resources, supervised and coordinated the work, and contributed to methodology, data curation, validation, visualisation, and manuscript preparation. Carlos Vivas-Rodríguez contributed to conceptual and methodological development, led software development, validation and visualisation, contributed to data curation, and led manuscript drafting. Olivier Dennler contributed to conceptual and methodological development, software, validation, supervision, visualisation, and manuscript revision. Colm Ryan contributed to conceptual and methodological development, software, resources, supervision, and manuscript revision. Siobhán McClean and David Gomez Matallanas contributed to conceptual development and critical manuscript review.

## Acknowledgements

The authors thank Dr Niamh Duggan and Ciarán Carey for testing *OrthoGather* with differential proteomics datasets during its development, and Dr Metin Yazar and Narod Kebabci for their technical assistance. In accordance with the journal policy on the use of generative AI, the authors acknowledge the use of ChatGPT, Codex and Claude Code to assist with language editing and debugging the software code. All outputs were reviewed, tested, and validated by the authors, who take full responsibility for the content.

## Data Availability

Source code, documentation and tutorials (*OrthoGather* v2.1.0, GPL-3.0-or-later) are available at GitHub (https://github.com/CarlosVivasR/OrthoGather) and Zenodo (https://doi.org/10.5281/zenodo.22738813).

## Supporting Information

**Supplementary Table S1**. Comparison of *OrthoGather* with representative tools for orthology inference, comparative genomics, and functional annotation.

**Supplementary File S1**. Analysis of the example dataset with OrthoGather.

**Supplementary File S2**. List of reference proteomes available through the Preselected Dataset input mode.

