## Supplementary File S1 - Example dataset analysis for "OrthoGather: a local platform for orthology-based proteome comparisons and Gene Ontology enrichment"

### Supplementary File S1. Analysis of an example dataset with *OrthoGather*

#### 1. Dataset used:

The example dataset was derived from Giddey *et al.* (2017). A temporal proteome dynamics study reveals the molecular basis of induced phenotypic resistance in *Mycobacterium smegmatis (*Msm*)* at sub-lethal rifampicin concentrations*. Scientific Reports*, 7:43858; PMID: 28262820.

Protein identifiers were obtained from the supplementary materials, which report quantitative proteomic profiles of Msm across three time points (30, 255 and 300 min) after exposure to sub-lethal rifampicin concentrations. Protein lists from all time points were pooled to create a dataset representing the overall response and were standardised to UniProt identifiers. Two subsets were extracted:

- *Foreground* (491 proteins): Proteins reported as differentially expressed at any of the time points.
- *Background* (3,181 proteins): The complete set of proteins detected in the study by liquid chromatography–tandem mass spectrometry (LC-MS/MS) and post MaxQuant processing with *M. smegmatis ATCC 700084* as a reference proteome.

These data were used to illustrate the complete analysis pipeline implemented in *OrthoGather* (Fig. S1) and were run using the *New Analysis* data input option. Tables containing the full list of the proteins are available at the end of this supplementary material and all *OrthoGather* output files from this analysis are available at https://github.com/CarlosVivasR/OrthoGather in the *example_analysis* folder.

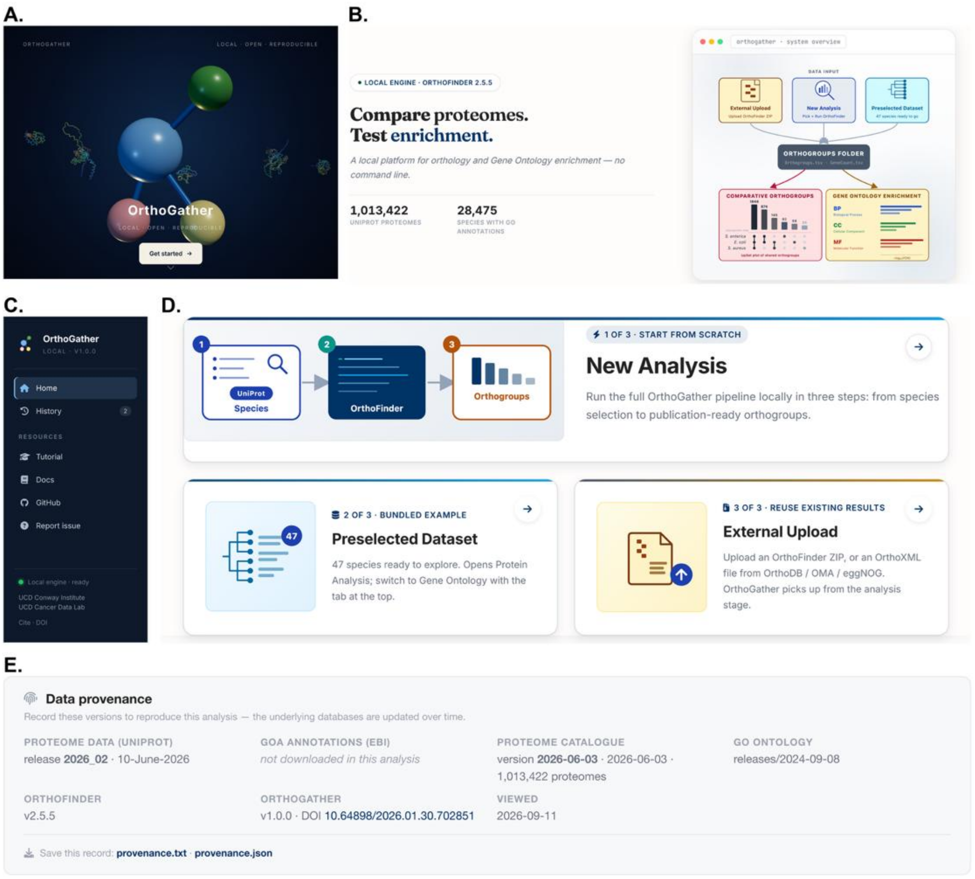


**Figure S1. Overview of the *OrthoGather* user interface. (A)** Starting view after opening *OrthoGather*. **(B)** Information panel highlighting the program version, and the available number of proteomes and GO annotations. **(C)** Navigation panel. **(D)** Data input options in *OrthoGather*. **(E)** Summary of the software and database versions used to generate the analysis, including the UniProt proteome release, UniProt proteome catalogue version, Gene Ontology release, OrthoFinder version, *OrthoGather* version, and analysis date. These details can be exported in text or JSON format to facilitate reproducibility and future reference.

We selected a subset of six bacterial species: *Acinetobacter baumannii* ATCC 19606, *Burkholderia cenocepacia* J2315, *Escherichia coli* K-12, *Mycobacteroides abscessus* ATCC 19977, *Mycolicibacterium smegmatis* ATCC 700084, and *Pseudomonas aeruginosa* PAO1 (Fig. S2). Msm is a rapidly growing non-tuberculous mycobacterium (NTM) widely used as a model organism in studies of mycobacterial disease and antibiotic resistance. Although generally considered non-pathogenic, rare case reports describe opportunistic infections, primarily in immunocompromised individuals. Its genome remains relatively poorly annotated, with many functional assignments inferred from other *Mycobacterium* species. *M. abscessus* (Mab*)*, a closely related NTM, is in contrast well recognised as an opportunistic pathogen and is increasingly viewed as transitioning towards a true pathogen. It is poorly annotated as well. *B. cenocepacia*, Mab, and *P. aeruginosa* represent typical cystic fibrosis (CF) pathogens, with *A. baumannii* also occasionally implicated in CF lung infections. The *E. coli* K-12 strain was included because it is among the most extensively annotated bacterial genomes and therefore contributes well-supported annotations to the orthogroups it shares with the other species. The selection of these specific species allows the user to assess whether proteins showing altered abundance in Msm following exposure to sub-lethal concentrations of rifampicin are species-specific or conserved in pathogenic mycobacteria and other CF-associated antibiotic-resistant pathogens, supporting hypothesis generation regarding potential rifampicin responses in those organisms.

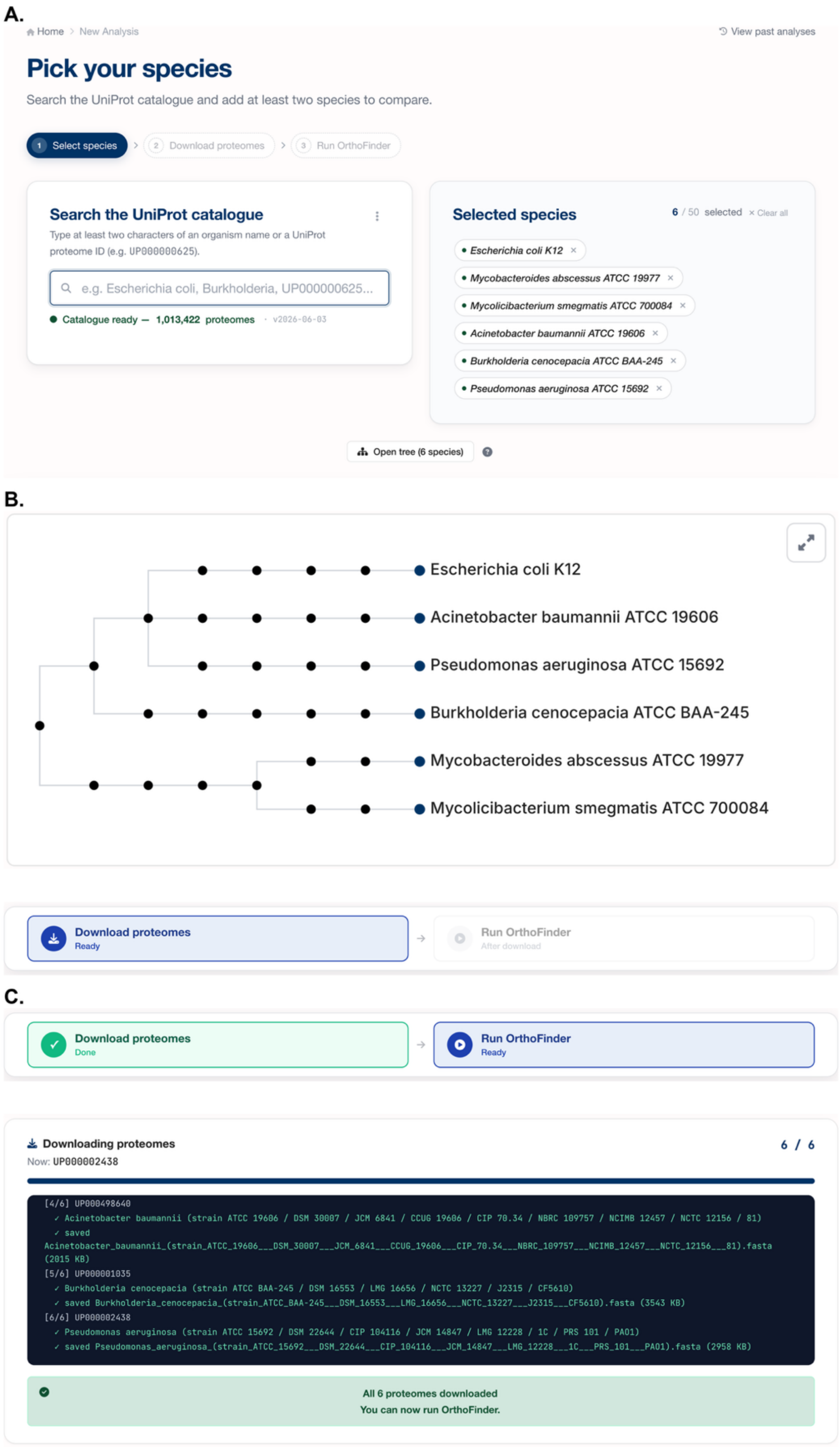


**Figure S2. Species selection interface for the *New Analysis* workflow in *OrthoGather.* (A)** Species selection panel used to define the organisms included in the analysis. The left panel provides access to the UniProt proteome catalogue, allowing species to be searched by name or UniProt proteome ID; the right panel shows the list of species currently selected by the user. **(B)** Taxonomic tree automatically generated from the selected species, in order to support species selection. The tree represents the NCBI taxonomic classification and is intended for visualisation of taxonomic relationships only, as it is not a sequence-based phylogeny. **(C)** Proteome download panel, showing the progress of proteome retrieval for all selected species, enabling users to monitor download status and overall program activity before OrthoFinder analysis is initiated.

#### 2. OrthoFinder outputs:

After downloading the proteomes, *OrthoFinder* can be executed. The analysis progress can be tracked by the user (Fig. S3A), and after the analysis is finished, summary statistics are generated (Fig. S3A), and the user can select one of the two analysis options (Fig. S3B). The four files (Table S2) can be downloaded as a ZIP archive from the run summary (Fig. S3A) or from any saved run in the Run History page.

**Table S2. OrthoFinder output files.**

| File | Description |
|---|---|
| *Orthogroups.tsv* | Lists all orthogroups and their corresponding UniProt identifiers for each species. |
| *Orthogroups.GeneCount.tsv* | Provides the number of genes (proteins) from each species within every orthogroup. |
| *Orthogroups_UnassignedGenes.tsv* | Contains genes that were not assigned to any orthogroup. |
| *Orthogroups.txt* | A compact text representation summarising all orthogroup compositions. |

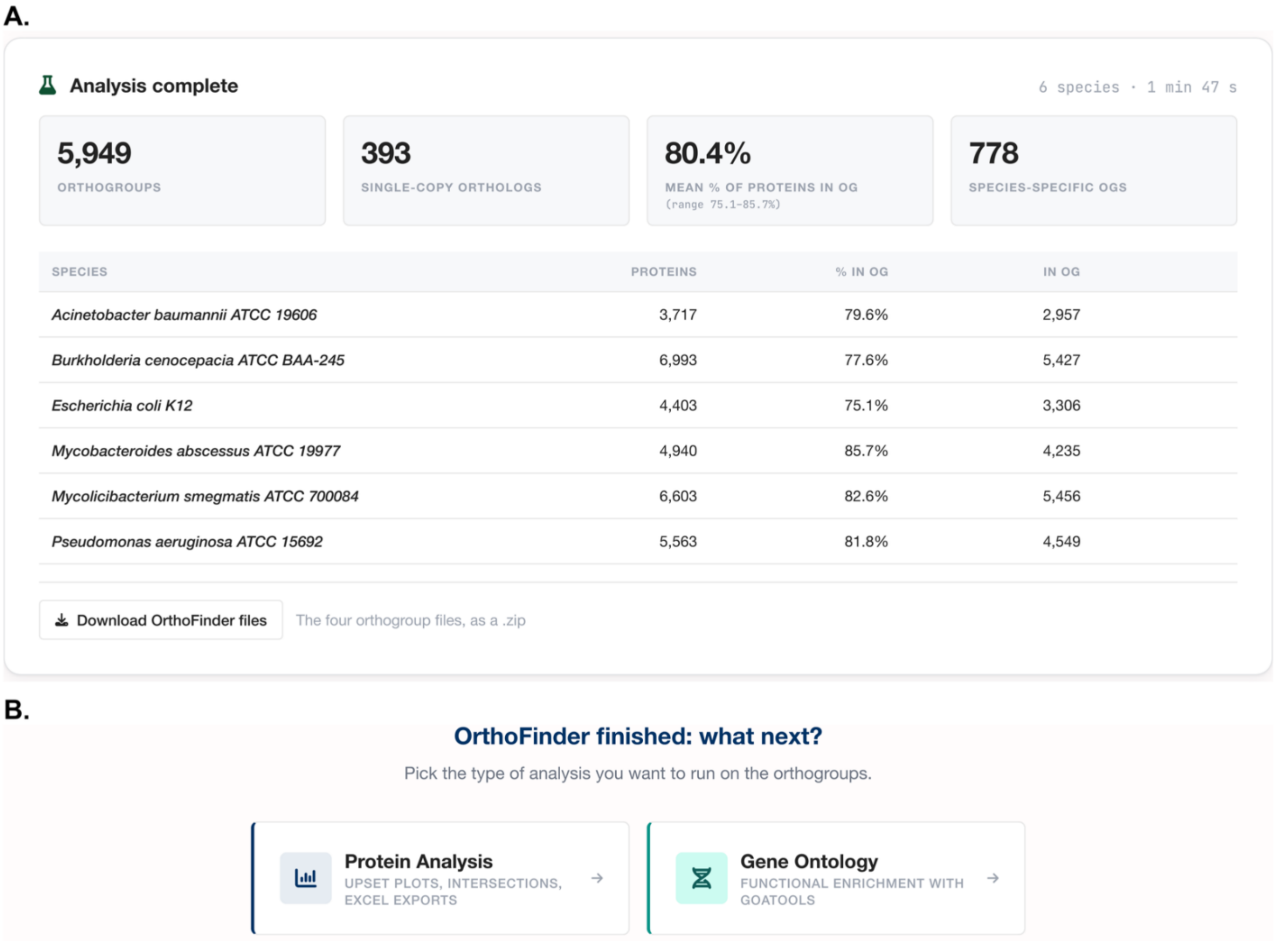


**Figure S3. OrthoFinder output summary in *OrthoGather*. (A)** Summary statistics generated after completion of the analysis. The upper panel reports the total number of orthogroups identified, the number of single-copy orthologues, the mean percentage of each species' proteins assigned to an orthogroup (with the range across species), and the number of species-specific orthogroups. The lower panel provides species-specific statistics, including the total number of proteins in each proteome, the percentage of proteins assigned to orthogroups, and the corresponding number of proteins assigned; the four OrthoFinder output files can be downloaded from this panel as a single .zip archive. **(B)** Analysis selection screen displayed after OrthoFinder completes, allowing the user to proceed with either comparative orthogroup analysis (Protein Analysis) or Gene Ontology enrichment analysis.

#### 3. Comparative Orthogroup Analysis

On entering the module, *OrthoGather* automatically generates global summary plots based on all species included in the preceding *OrthoFinder* analysis (Fig. S4A). These provide an overview of the distribution of proteins per orthogroup, and the number of species represented within each orthogroup. The user can then select a subset of the analysed species for more detailed comparative analysis (Fig. S4B). The selected subset is used to generate interactive UpSet plots showing orthogroup and protein intersections among those species (Fig. S4C–D). In this example, the initial analysis contained all six bacterial species, while Msm, Mab and *E. coli* were selected for the subsequent UpSet analysis to simplify visualisation and interpretation of species intersections.

This step generated two UpSet plots illustrating the distribution of orthogroups (Fig. S4C) and proteins (Fig. S4D) among the selected species. The complete underlying data for these visualisations are provided in the *example_analysis/2_comparative_analysis* folder on GitHub, as one Excel file per UpSet plot. Each file contains detailed protein composition for each orthogroup corresponding to the species intersections shown in the UpSet plots. The first worksheet summarises the content of the file and the README tab contains information about the analysis such as date and time of generation, species analysed and the number of species combinations in the plot. Each of the following worksheets represents a specific intersection combination corresponding to a unique subset of species that share orthogroups and proteins. Within each worksheet, rows correspond to orthogroups, while columns are defined as: *Orthogroup ID*: Identifier of the orthogroup as reported by OrthoFinder, unique for each separate analysis. *Subsequent columns*: UniProt accession codes of the proteins from each species present in the orthogroup. *Empty cells*: Indicate that the corresponding species does not contribute proteins to that orthogroup in the given intersection.

The UpSet plots generated for this example dataset show that 447 orthogroups encompassing 817 proteins are unique to Msm*,* whereas 1,024 orthogroups comprising 4,599 proteins are shared by all three selected species. Finally, Mab and Msm, as closely related species, share 1,699 orthogroups encompassing 4,709 proteins (Fig. S4C–D). It is important to note that one orthogroup may contain multiple proteins from a species or not contain any for a given species.

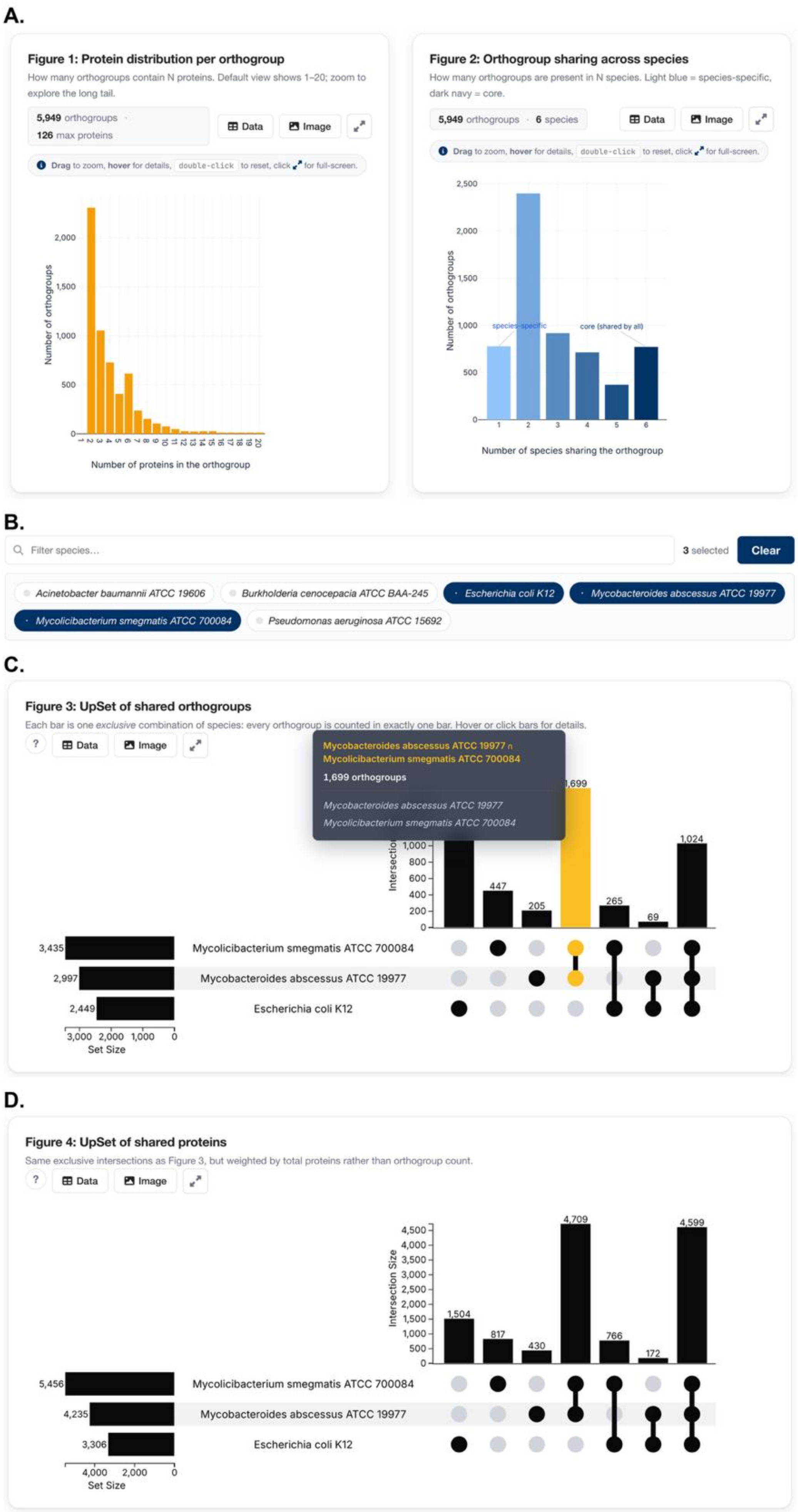


**Figure S4. Comparative orthogroup analysis in *OrthoGather.*** **(A)** Global summary plots showing the distribution of proteins per orthogroup (left) and the distribution of orthogroups according to the number of species in which they are present (right). **(B)** Species selection panel used to define the subset of species included in the comparative analysis and subsequent UpSet plots. In this example, the analysis was restricted to Msm, Mab and *E. coli*. **(C–D) Interactive UpSet** plots illustrating orthogroup and protein overlap among the selected species. **(C)** UpSet plot showing the number of orthogroups unique to or shared among the selected species; hovering over bars or intersections highlights the selected combination in yellow and displays detailed information. **(D)** UpSet plot showing the corresponding distribution of proteins within these orthogroup intersections. In both UpSet plots, the left horizontal bar chart displays the total number of orthogroups, or proteins, associated with each species, whereas the upper vertical bar chart shows the size of each species intersection defined by the matrix below.

#### 4. Filtered Orthogroup Analysis

This module allows the user to filter the orthogroups with a user-defined protein identifier list (Fig. S5). We have used the 491 Msm proteins mentioned above, of which 467 were mapped to 430 orthogroups. The remaining 24 IDs were not assigned to any orthogroup by OrthoFinder and were therefore not included in the subsequent UpSet plots. These proteins did not group with any homologous proteins within the analysed dataset, including proteins from Msm itself. This is distinct from Msm-specific orthogroups, which contain proteins found only in Msm but still include two or more proteins grouped together based on sequence similarity.

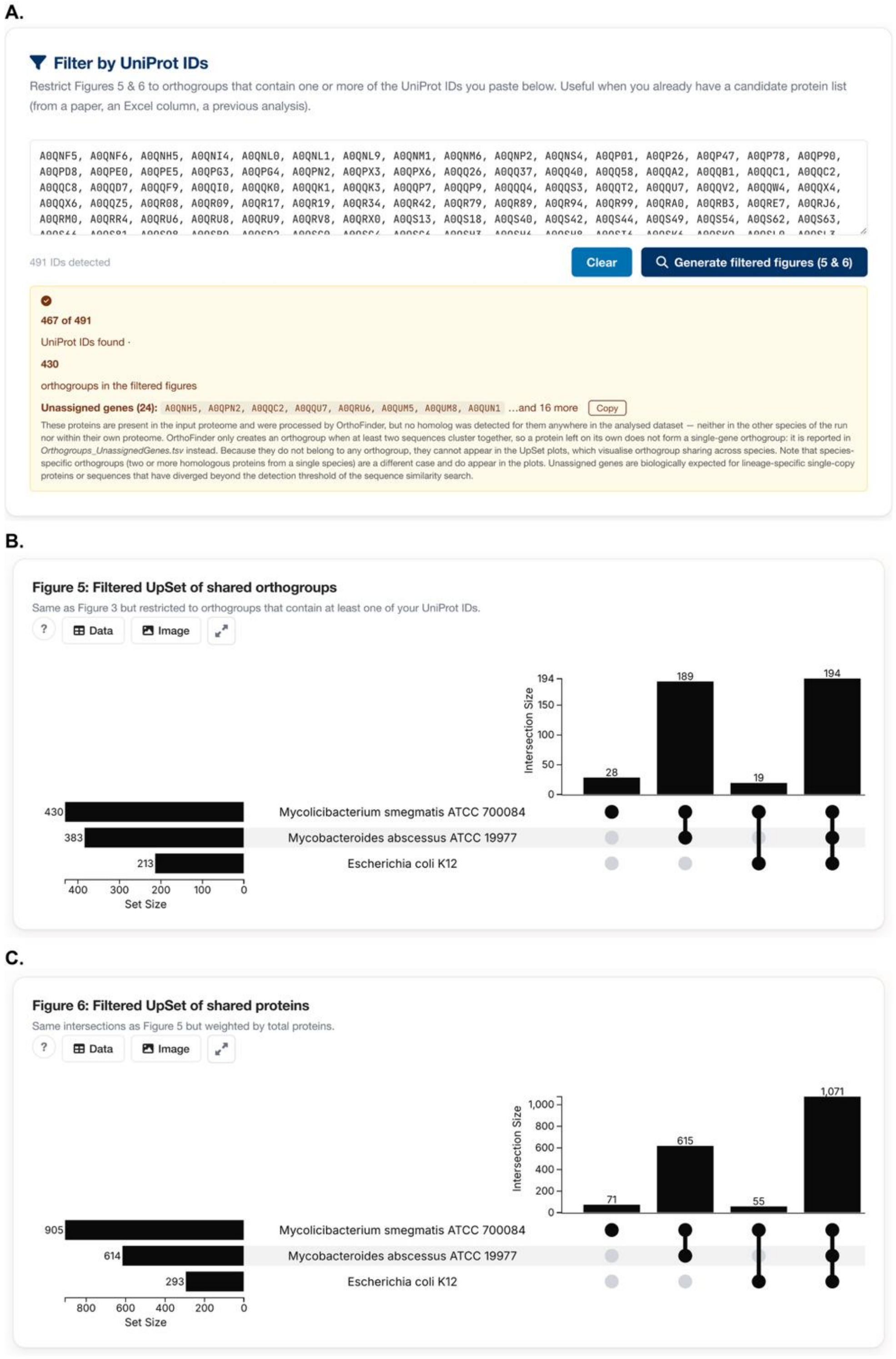


**Figure S5. Defining a protein subset in the filtered orthogroup analysis in *OrthoGather.*** **(A)** Option of filtering the UpSet plots by a defined list of UniProt IDs. (B–C) UpSet plots generated following subsequent filtering of the example dataset using the foreground protein set, retaining only orthogroups containing at least one selected protein. **(B)** UpSet plot showing the number of retained orthogroups in each species and their intersections. **(C)** UpSet plot showing the number of proteins within the corresponding retained orthogroup intersections. In all UpSet plots, the left bar chart shows the total number of orthogroups, or proteins, associated with each species, while the upper bar chart shows the size of each exclusive intersection defined by the species combinations indicated below.

These proteins were mapped onto the existing orthogroups obtained from the global analysis, and another two UpSet plots were generated in this step, specifically reflecting the distribution of retained orthogroups and all proteins contained within them across the three selected bacterial species. The first visualises orthogroup distribution (Fig. S5B), and the second one visualises protein distribution (Fig. S5C). As in the previous step, an accompanying Excel file is generated, containing the detailed corresponding data (GitHub *example_analysis/2_comparative_analysis* folder)

The two UpSet plots generated in this step reveal 28 orthogroups unique to Msm, comprising 71 proteins in total. In the Excel data sheet, a corresponding tab lists this information, including the protein IDs (Table S3). Of the 71 proteins, 35 belonged to the defined Msm foreground set and were distributed across all 28 Msm-specific orthogroups. Five orthogroups contained more than one foreground protein, accounting for 12 of the 35 proteins, while the remaining 23 orthogroups each contained a single foreground protein. Thirteen of the 28 Msm-specific orthogroups consisted exclusively of proteins from the foreground set, whereas the remaining 15 also contained Msm-specific proteins not included in the foreground set. Users can refer back to UniProt to obtain additional information on the proteins, including their names and predicted or annotated functions. For example, proteins within OG0000261 are annotated as cutinases, proteins within OG0000443 as putative S-adenosyl-L-methionine-dependent methyltransferases, and proteins within OG0001856 as glycerol and propanediol dehydratases.

**Table S3. Filtered orthogroups unique to Msm**, as found in *orthogather_figure5-upset-orthogroups-filtered_3species_430OGs-filtered.xlsx* (*example_analysis/2_comparative_analysis*)**.** Bold: proteins defined by us (Table S5). For each orthogroup only the proteins present in the filtering list are shown, so a single identifier in this table does not indicate a single-protein orthogroup.

| Orthogroup ID | Mycolicibacterium_smegmatis_ (strain_ATCC_700084___mc(2)155) |
|---|---|
| OG0000261 | tr\|A0QNX2\|A0QNX2_MYCS2, tr\|A0QRN8\|A0QRN8_MYCS2, tr\|A0QSA0\|A0QSA0_MYCS2, tr\|A0QSM1\|A0QSM1_MYCS2, **tr\|A0QSM2\|A0QSM2_MYCS2**, **tr\|A0QSM3\|A0QSM3_MYCS2**, tr\|A0QU66\|A0QU66_MYCS2, tr\|A0QV75\|A0QV75_MYCS2, tr\|A0R0Q0\|A0R0Q0_MYCS2, tr\|A0R4L6\|A0R4L6_MYCS2, tr\|A0R5Y1\|A0R5Y1_MYCS2 |
| OG0000443 | **sp\|A0QNM1\|Y093_MYCS2**, sp\|A0QNM2\|Y095_MYCS2, **sp\|A0QQ37\|Y614_MYCS2**, **sp\|A0QSH3\|Y1479_MYCS2**, sp\|A0QSH4\|Y1480_MYCS2, sp\|A0QSH5\|Y1481_MYCS2, **sp\|A0QSH6\|Y1482_MYCS2**, sp\|A0QTL6\|Y1888_MYCS2, tr\|A0QQ31\|A0QQ31_MYCS2 |
| OG0000550 | **tr\|A0R0I3\|A0R0I3_MYCS2**, tr\|A0R239\|A0R239_MYCS2 |
| OG0000741 | **tr\|A0R003\|A0R003_MYCS2** |
| OG0001226 | **tr\|A0R2M2\|A0R2M2_MYCS2** |
| OG0001608 | **tr\|A0R3U1\|A0R3U1_MYCS2**, tr\|A0R5P4\|A0R5P4_MYCS2 |
| OG0001756 | tr\|A0QSK1\|A0QSK1_MYCS2, **tr\|A0QU64\|A0QU64_MYCS2** |
| OG0001856 | tr\|A0QPS1\|A0QPS1_MYCS2, **tr\|A0QSP1\|A0QSP1_MYCS2**, **tr\|A0QSP2\|A0QSP2_MYCS2**, tr\|A0QSP3\|A0QSP3_MYCS2, tr\|A0R5V1\|A0R5V1_MYCS2 |
| OG0002291 | **tr\|A0QPX3\|A0QPX3_MYCS2**, tr\|A0QVN3\|A0QVN3_MYCS2 |
| OG0002362 | **tr\|A0R6E9\|A0R6E9_MYCS2**, tr\|A0R6T8\|A0R6T8_MYCS2 |
| OG0002562 | tr\|A0QPC7\|A0QPC7_MYCS2, tr\|A0QW96\|A0QW96_MYCS2, **tr\|A0R1J0\|A0R1J0_MYCS2**, tr\|A0R2R6\|A0R2R6_MYCS2 |
| OG0002571 | tr\|A0QS70\|A0QS70_MYCS2, tr\|A0R181\|A0R181_MYCS2, tr\|A0R505\|A0R505_MYCS2, **tr\|A0R771\|A0R771_MYCS2** |

| Orthogroup ID | Mycolicibacterium_smegmatis_ (strain_ATCC_700084___mc(2)155) |
|---|---|
| OG0002578 | tr\|A0QZF5\|A0QZF5_MYCS2, tr\|A0QZF6\|A0QZF6_MYCS2, **tr\|A0R1A9\|A0R1A9_MYCS2** |
| OG0002967 | **tr\|A0QWH3\|A0QWH3_MYCS2** |
| OG0003091 | **tr\|A0QRR4\|A0QRR4_MYCS2** |
| OG0003188 | tr\|A0QT82\|A0QT82_MYCS2, **tr\|A0R3W0\|A0R3W0_MYCS2** |
| OG0003237 | **tr\|A0QX61\|A0QX61_MYCS2** |
| OG0003599 | tr\|A0QNK0\|A0QNK0_MYCS2, tr\|A0QW60\|A0QW60_MYCS2, **tr\|A0R130\|A0R130_MYCS2** |
| OG0003624 | tr\|A0R1J8\|A0R1J8_MYCS2, tr\|A0R517\|A0R517_MYCS2, **tr\|A0R6P9\|A0R6P9_MYCS2** |
| OG0003664 | **sp\|A0R480\|KDC_MYCS2** |
| OG0003938 | **tr\|A0QWM9\|A0QWM9_MYCS2** |
| OG0003975 | **tr\|A0R189\|A0R189_MYCS2** |
| OG0004215 | **tr\|A0QWI4\|A0QWI4_MYCS2** |
| OG0004266 | **tr\|A0R1H5\|A0R1H5_MYCS2** |
| OG0005759 | **tr\|A0QZ85\|A0QZ85_MYCS2**, **tr\|A0R1C8\|A0R1C8_MYCS2** |
| OG0005763 | **tr\|A0QZ98\|A0QZ98_MYCS2**, **tr\|A0R2V0\|A0R2V0_MYCS2** |
| OG0005792 | tr\|A0QRE5\|A0QRE5_MYCS2, **tr\|A0QRE7\|A0QRE7_MYCS2** |
| OG0005800 | **tr\|A0QS13\|A0QS13_MYCS2** |

#### *5.* Gene Ontology Enrichment Analysis

A separate *Gene Ontology* module in *OrthoGathe*r enables functional interpretation of inferred orthogroups, both before and after filtering, provided that the analysed species have corresponding annotations available in the Gene Ontology Annotation (GOA) repository. This module supports GO enrichment analysis to identify functional categories that are over-represented within user-defined protein sets, allowing biological processes, molecular functions, or cellular components associated with specific experimental conditions to be highlighted. It is summarised in an Excel file (GitHub *example_analysis/3_go_annotation* folder; Table S4)

**Table S4. Summary of GO term analysis.** Worksheets document the specific steps of the GO analysis process (*orthogather_GOA-orthogroup-annotation_6species.xlsx*, *example_analysis/3_go_annotation*).

| Sheet name | Description |
|---|---|
| Meta | General analysis metadata: total number of orthogroups (*n_orthogroups*), number of species columns (*n_species_cols*), annotation coverage metrics (*rows_nonzero_annotation* and *rows_zero_annotation*). |
| Initial Groups | List of orthogroups obtained directly from *OrthoFinder* before any filtering or annotation-based selection. |
| Filtered Groups | All orthogroups containing at least one annotated protein. |
| Removed Groups | Orthogroups in which no protein carries a GO annotation, and which therefore contribute nothing to enrichment. |
| Groups of Interest | All orthogroups, restricted to the species columns that have a GOA file. This is the sheet the enrichment reads. |
| Species & GOA Map | Mapping between each analysed species and the corresponding GOA file. |

Firstly, prior to GO enrichment analysis, *OrthoGather* verifies the availability of GOA data for each analysed species and identifies those with corresponding annotations for downstream analysis (Fig. S6A). Where a GOA file cannot be matched automatically, the match can be corrected manually by providing a Taxon ID. If no valid GOA file is available, proteins from that species do not contribute GO annotations to the enrichment analysis, but they remain part of the orthogroup structure. Proteins without assigned GO terms may still be present but do not contribute to enrichment statistics.

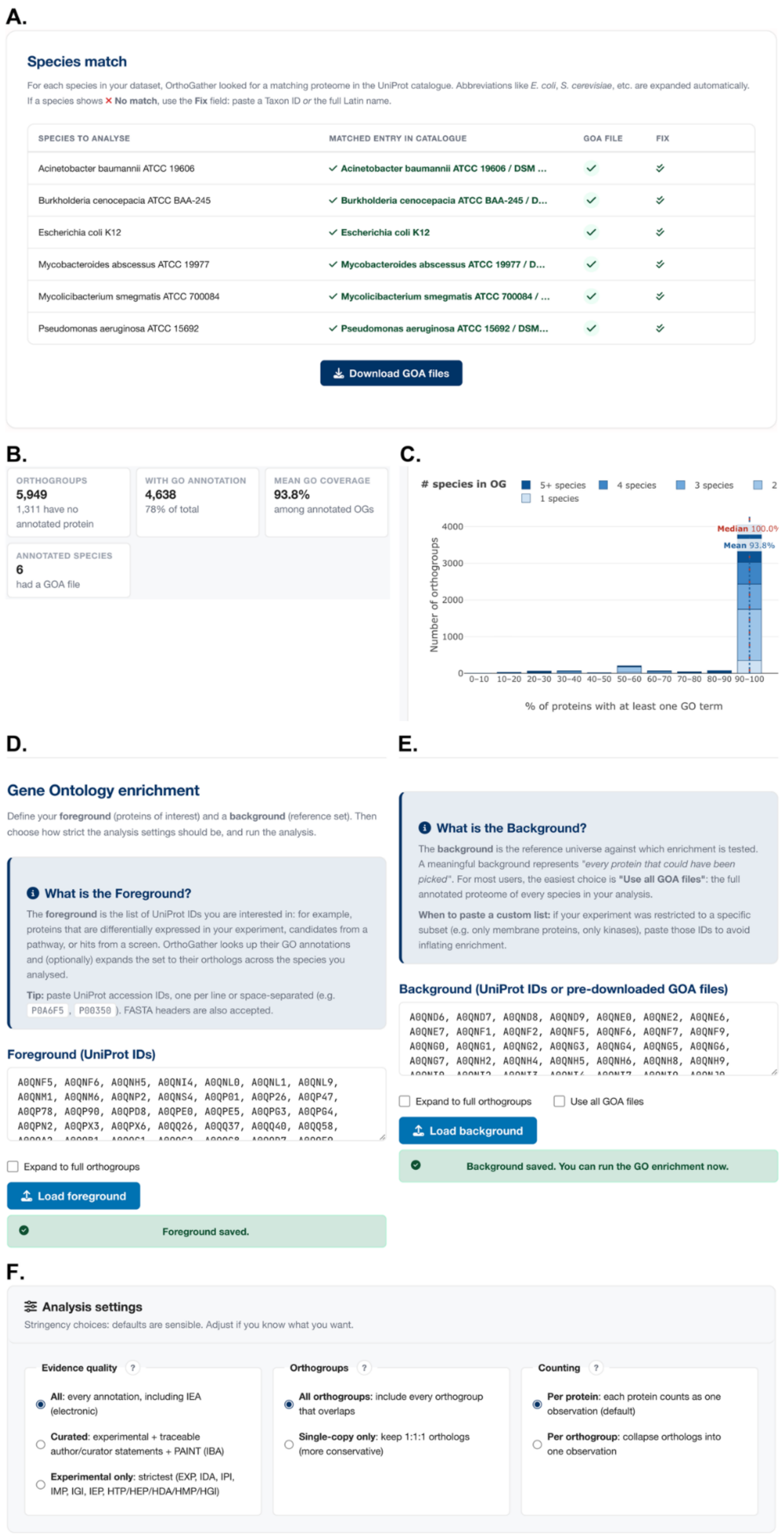


**Figure S6. Verification of Gene Ontology Annotation (GOA) file availability and Gene Ontology enrichment analysis setup in *OrthoGather*.** **(A)** Automatic matching of the selected species to GOA files available in the UniProt GOA repository. If no matching GOA file is identified, the user can manually specify the corresponding NCBI Taxonomy ID to search for an alternative file. Species without an available GOA file remain included in the orthogroup analysis but do not contribute Gene Ontology annotations to the enrichment analysis. **(B)** Annotation overview reporting the total number of orthogroups, the number containing at least one protein with a GO term, their mean annotation coverage, and the number of species with an available GOA file. **(C)** Distribution of Gene Ontology annotation coverage across orthogroups, defined as the percentage of proteins within an orthogroup with at least one GO term; orthogroups are stacked by the number of species they contain. **(D–F) Configuration** of Gene Ontology enrichment analysis. **(D)** Definition of the foreground protein set (proteins of interest), with the option to expand it to orthologous proteins from the selected species. **(E)** Definition of the background protein set (reference dataset), with options to include all orthologous proteins or all GOA files. **(F)** Gene Ontology enrichment settings, including annotation evidence quality (all annotations, non-IEA annotations only, or experimental evidence only), all or only single-copy orthogroups, and counting strategy, with each orthologous protein treated as an individual observation (per protein) or all orthologues within an orthogroup collapsed into a single observation (per orthogroup).

After the GOA files are downloaded, *OrthoGather* provides an annotation overview summarising GO annotation coverage across the dataset (Fig. S6B). This reports the total number of orthogroups, the number lacking annotation, the number and proportion containing at least one annotated protein, the mean annotation percentage among annotated orthogroups, and the number of species for which a GOA file was available. GOA files were available for all six species. In the example dataset, 4,638 of the 5,949 orthogroups (78.0%) contained at least one annotated protein, with a mean annotation percentage of 93.8% among them; the remaining 1,311 orthogroups contain no annotated protein and therefore contribute nothing to the enrichment. A more detailed view of per-orthogroup annotation coverage is provided by the annotation distribution chart (Fig. S6C).

Further, in the enrichment phase, the user can perform a GO enrichment analysis for a defined subset of proteins, based on GO terms assigned to proteins within the selected orthogroups from species with available GOA files. The foreground (proteins of interest) and the background against which it is tested must be defined (Fig. S6D–F). For this example dataset analysis 491 differentially expressed proteins returned a single significant term (protein maturation), thus, they were therefore split into two separate foregrounds: 185 increased in abundance and 306 decreased in abundance, and analysed independently against a common background of the 3,181 Msm proteins detected by mass spectrometry. All evidence codes, per-protein counting and all orthogroups were used, and the foreground was analysed as pasted, without expansion to the full orthogroups, so that the enrichment describes the Msm proteins reported by the original study. Of the background, 2,470 proteins carry at least one GO term and form the population for the test; 162 of the 185 increased-in-abundance and 250 of the 306 decreased-in-abundance foreground proteins are used. *OrthoGather* restricts the foreground to proteins that are both GO-annotated and present in the annotated background. Therefore, although 251 of the 306 decreased-in-abundance identifiers carry a GO annotation, only 250 are present in the background and included in the enrichment test.

To determine whether a GO term is over-represented in the foreground relative to the background, a one-sided Fisher's exact test is applied (GOATOOLS v1.6.4, go-basic.obo release 2026-07-26). Annotations are propagated to parent terms along is_a and part_of relationships, annotations carrying the NOT qualifier are discarded, and terms below the selected minimum GO hierarchy depth, or annotating fewer than 10 or more than 500 proteins of the reference universe, are removed before correction. Multiple testing is then corrected with the Benjamini–Hochberg false discovery rate (FDR). As in PANTHER, the three namespaces share a single annotated reference universe; the correction is computed once across all three. *OrthoGather* generates interactive, adjustable figures, which allow the user to display either all categories on one bar chart or separate them based on GO component, displaying GO term enrichment for Biological Process, Molecular Function and Cellular Component (Fig. S7), and an accompanying data file summarising the statistics and results (GitHub *example_analysis/4_go_enrichment* folder. The user can specify the FDR threshold, the minimum GO hierarchy depth, the maximum number of terms displayed per namespace, and a separate display threshold that draws near-significant terms as hollow bars so that they remain visible without being reported as findings.

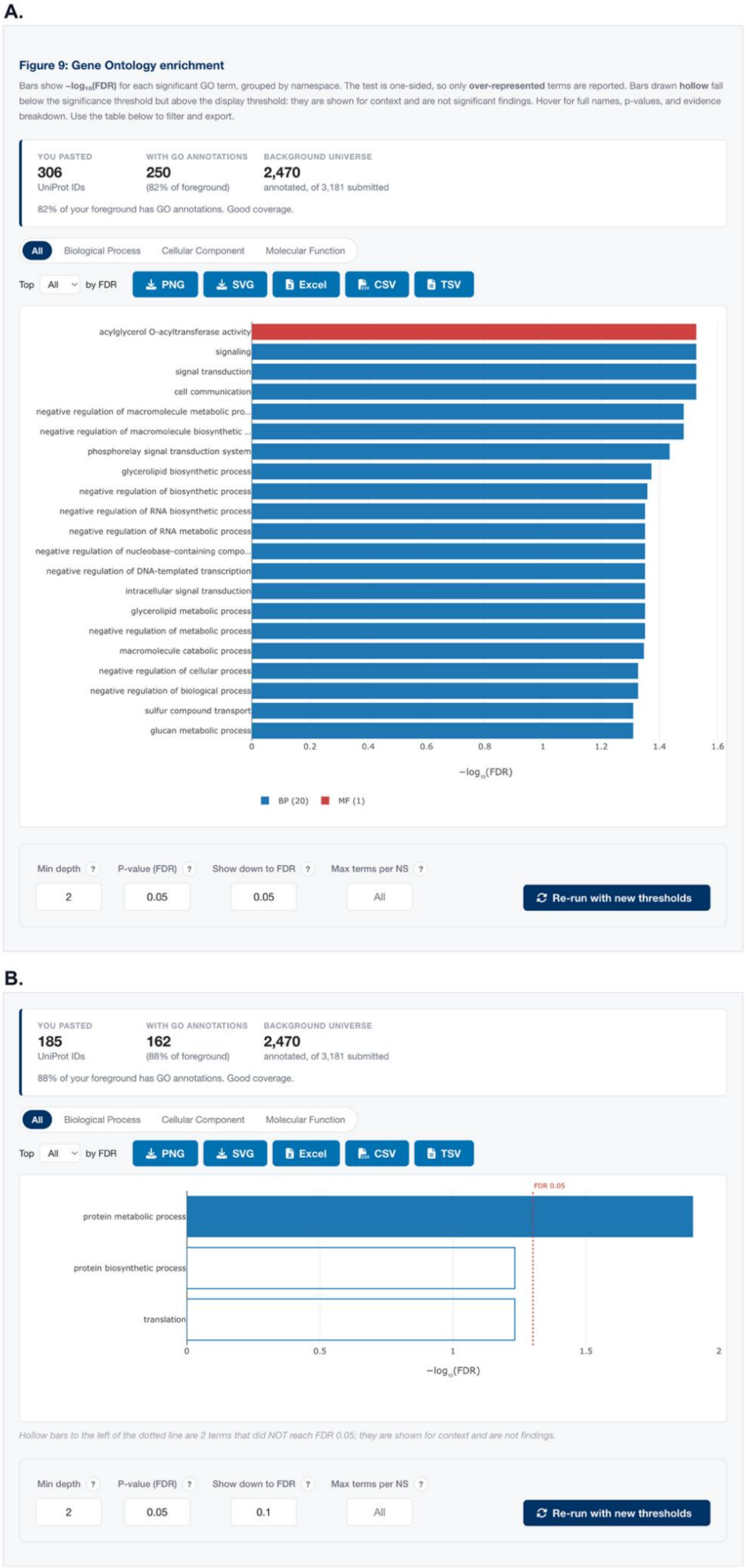


**Figure S7. Gene Ontology enrichment results for proteins increased and decreased in abundance.** Foreground and background sets are defined in Table S5. GO enrichment was performed using a false discovery rate (FDR) threshold of 0.05, a minimum GO hierarchy depth of 2, and no limit on the number of terms displayed. Results are grouped by the three Gene Ontology namespaces: Biological Process (BP, blue), Molecular Function (MF, red), and Cellular Component (CC, green). Each horizontal bar represents an enriched GO term, with bar length corresponding to $-\log_{10}$(FDR). The upper summary panel reports the foreground size, the number of GO-annotated foreground proteins included in the analysis, and the annotated background size. **(A)** Analysis of the 306 proteins decreased in abundance, of which 250 (82%) were annotated and present in the background, identified 21 enriched terms: 20 Biological Process terms and one Molecular Function term. **(B)** Analysis of the 185 proteins increased in abundance, of which 162 (88%) were annotated, identified a single enriched term, protein metabolic process. **Hollow bars:** terms with FDR $\geq 0.05$ and $< 0.10$, shown for context only. **Dotted line:** FDR = 0.05 significance threshold. Analysis settings are displayed below each panel.

With the FDR threshold at 0.05, the minimum GO hierarchy depth at two and no limit on the number of terms displayed, the decreased-in-abundance foreground returned 21 enriched terms, 20 Biological Process and one Molecular Function (Fig. S7A; Fig. S8A), whereas the increased-in-abundance foreground returned a single enriched term (Fig. S7B; Fig. S8B). No Cellular Component terms were significantly enriched. The decreased-in-abundance foreground showed enrichment of Biological Process terms associated with negative regulation of transcription, biosynthesis and metabolism, signal transduction, glycerolipid metabolism and biosynthesis, macromolecule catabolism, glucan metabolism, and sulfur compound transport. Acylglycerol O-acyltransferase activity was the only significantly enriched Molecular Function term. Raising the minimum hierarchy depth to five retained 14 of the 21 terms and additionally identified phospholipid metabolic process as significantly enriched.

Among the proteins decreased in abundance, the enrichment of signal transduction processes supports the involvement of two-component regulatory systems in coordinating the cellular response to rifampicin stress reported in the original study, and the recurrence of negative-regulation terms reflects a single group of 13 regulatory proteins, 11 of which carry the transcription-related terms. The enrichment of glycerolipid metabolic and biosynthetic processes, together with acylglycerol O-acyltransferase activity, is consistent with the remodelling of lipid metabolism and the cell envelope reported in the original study during adaptation to rifampicin exposure. Among proteins increased in abundance, only one term was significantly enriched, while translation and protein biosynthetic process did not reach the significance threshold (FDR = 0.06), consistent with the transcriptional and translational response to rifampicin-mediated inhibition of transcription described in the original study. As 98.3% of the GO annotations available for Msm are inferred electronically (IEA), the enriched terms primarily reflect the functional annotations associated with these proteins rather than experimentally validated processes in Msm.

Overall, this analysis showed that *OrthoGather's* orthogroup-focused enrichment framework recapitulates major biological responses reported by the original proteomic study while enabling interactive exploration of the underlying orthogroups. Users can examine the distribution of orthologous proteins and their functional annotations, identify the presence or absence of orthologues across selected species, and use this comparative information to formulate hypotheses regarding protein conservation, divergence, and biological roles.

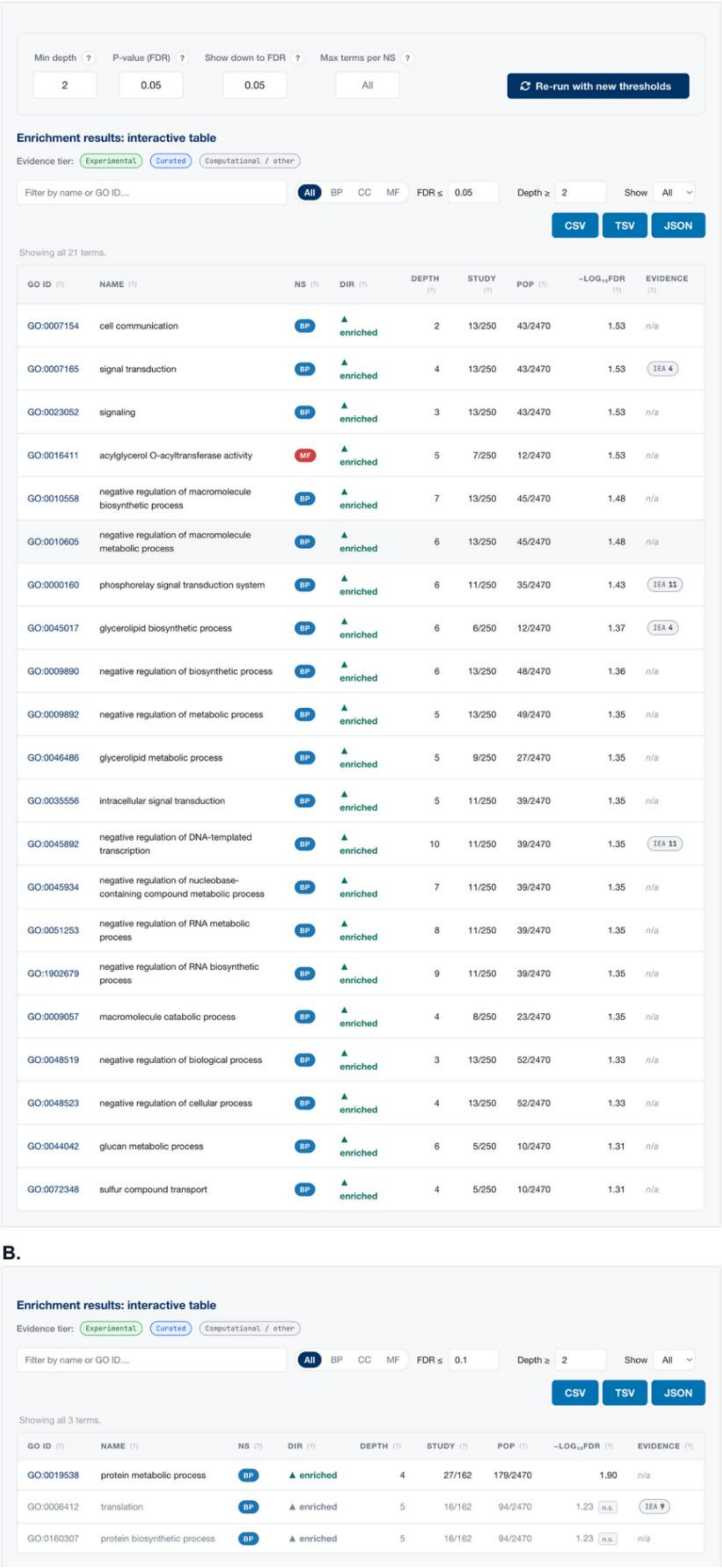


**Figure S8. Interactive table of enriched Gene Ontology terms generated by *OrthoGather*. (A)** Results for proteins decreased in abundance, comprising 21 enriched terms. The table reports the GO identifier, term name, ontology namespace (Biological Process (BP), Cellular Component (CC), or Molecular Function (MF)), enrichment direction, hierarchy depth, the number of foreground proteins or orthogroups associated with each term (Study), the corresponding number in the annotated background (POP), $-\log_{10}$(FDR), and GO annotation evidence codes (Evidence; e.g. IDA: Inferred from Direct Assay; ISS: Inferred from Sequence or Structural Similarity; IEA: Inferred from Electronic Annotation). Results can be filtered by term name or GO identifier, ontology namespace, evidence type, FDR threshold, and minimum hierarchy depth, and exported as CSV, TSV, or JSON. **(B)** Results for proteins increased in abundance, with terms at FDR ≤ 0.10 displayed. Terms with FDR ≥ 0.05 are marked in grey and labelled n.s. Analysis settings are shown above each table.

**Table S5. Foreground and background IDs used for the example analysis.**

| **Foreground (n = 491)** |
|---|
| **Foreground - down-regulated (n = 306)**<br>A0QNF5, A0QNF6, A0QNH5, A0QNL9, A0QNP2, A0QNS4, A0QP01, A0QP26, A0QP47, A0QP78, A0QP90, A0QPD8, A0QPE0, A0QPN2, A0QPX3, A0QPX6, A0QQ58, A0QQA2, A0QQB1, A0QQC2, A0QQF9, A0QQI0, A0QQK1, A0QQK3, A0QQP7, A0QQP9, A0QQS3, A0QQT2, A0QQU7, A0QQZ5, A0QR08, A0QR09, A0QR17, A0QR34, A0QR42, A0QR94, A0QRA0, A0QRM0, A0QRU6, A0QRU8, A0QRU9, A0QS13, A0QS81, A0QSB9, A0QSG4, A0QSI6, A0QSK6, A0QSK9, A0QSL0, A0QSP1, A0QSP2, A0QSQ6, A0QSV0, A0QT07, A0QT09, A0QT13, A0QT17, A0QT20, A0QT33, A0QT70, A0QTD8, A0QTE3, A0QTF7, A0QTG8, A0QTK2, A0QTK6, A0QTL7, A0QTR3, A0QTS2, A0QU18, A0QU43, A0QU51, A0QU56, A0QU64, A0QUA6, A0QUI9, A0QUM5, A0QUM6, A0QUM7, A0QUM8, A0QUN1, A0QUN2, A0QUN3, A0QUN5, A0QUN7, A0QUN8, A0QUN9, A0QUP0, A0QUW6, A0QUW7, A0QUZ3, A0QUZ6, A0QV14, A0QV18, A0QV25, A0QV52, A0QVJ7, A0QVL0, A0QVL4, A0QVL5, A0QVL6, A0QVP7, A0QVQ8, A0QVS0, A0QVT0, A0QVT1, A0QVY5, A0QVY8, A0QVY9, A0QVZ3, A0QW04, A0QW19, A0QW21, A0QW22, A0QW25, A0QWG0, A0QWG5, A0QWG6, A0QWH2, A0QWH3, A0QWH5, A0QWI4, A0QWR3, A0QWR9, A0QWU1, A0QWU2, A0QWV0, A0QWV7, A0QWX6, A0QWY3, A0QX55, A0QX61, A0QX62, A0QX75, A0QX76, A0QX77, A0QXA6, A0QXA7, A0QXA8, A0QXB5, A0QXB9, A0QXC6, A0QXD2, A0QXK4, A0QXP3, A0QXX7, A0QXY1, A0QXZ3, A0QY08, A0QY24, A0QY91, A0QY95, A0QYA9, A0QYD3, A0QYE3, A0QYE5, A0QYG2, A0QYQ2, A0QYQ8, A0QYU8, A0QYW5, A0QZ16, A0QZ42, A0QZ46, A0QZ54, A0QZ56, A0QZ60, A0QZ83, A0QZ85, A0QZ86, A0QZ91, A0QZ95, A0QZ98, A0QZA1, A0QZA2, A0QZA6, A0R003, A0R012, A0R031, A0R037, A0R045, A0R051, A0R090, A0R091, A0R097, A0R0B2, A0R0B4, A0R0B5, A0R0B7, A0R0I3, A0R0Q5, A0R0Q9, A0R0R3, A0R0R8, A0R0S9, A0R0T9, A0R0W2, A0R0W4, A0R0W7, A0R0W9, A0R0Y4, A0R170, A0R171, A0R196, A0R1C2, A0R1C6, A0R1C7, A0R1C8, A0R1D5, A0R1E5, A0R1H5, A0R1J0, A0R1X5, A0R1X8, A0R1Y7, A0R1Y8, A0R217, A0R266, A0R268, A0R2B2, A0R2C4, A0R2D2, A0R2E1, A0R2E2, A0R2J4, A0R2P0, A0R2V0, A0R2V2, A0R2V3, A0R2V4, A0R2V6, A0R2V8, A0R2V9, A0R2W6, A0R2W7, A0R2Y3, A0R316, A0R325, A0R368, A0R3A6, A0R3C6, A0R3D7, A0R3F3, A0R3H9, A0R3I6, A0R3I7, A0R3K5, A0R3Q0, A0R3Q2, A0R3R3, A0R3S7, A0R3Y4, A0R403, A0R407, A0R408, A0R420, A0R430, A0R443, A0R461, A0R471, A0R478, A0R480, A0R4B3, A0R4E0, A0R4F7, A0R4H6, A0R4L1, A0R4N3, A0R4Q1, A0R518, A0R565, A0R566, A0R588, A0R592, A0R595, A0R5C0, A0R5E5, A0R5I4, A0R5I5, A0R5K2, A0R5M8, A0R5N0, A0R5Q8, A0R5R4, A0R5R9, A0R5T7, A0R5T8, A0R5W8, A0R5Y9, A0R610, A0R624, A0R628, A0R6C7, A0R6D2, A0R6P9, A0R716, A0R722, A0R726, A0R727, A0R771, O52200, P0CG99, P0CH00, P94968, Q50441, Q9F868<br><br>**Foreground - up-regulated (n = 185)**<br>A0QNI4, A0QNL0, A0QNL1, A0QNM1, A0QNM6, A0QPE5, A0QPG3, A0QPG4, A0QQ26, A0QQ37, A0QQ40, A0QQC1, A0QQC8, A0QQD7, A0QQK0, A0QQQ4, A0QQV2, A0QQW4, A0QQX4, A0QQX6, A0QR19, A0QR79, A0QR89, A0QR99, A0QRB3, A0QRE7, A0QRJ6, A0QRR4, A0QRV8, A0QRX0, A0QS18, A0QS40, A0QS42, A0QS44, A0QS49, A0QS54, A0QS62, A0QS63, A0QS66, A0QS98, A0QSD2, A0QSG0, A0QSG6, A0QSH3, A0QSH6, A0QSH8, A0QSL3, A0QSL8, A0QSM2, A0QSM3, A0QSR8, A0QSS4, A0QT04, A0QT11, A0QTG7, A0QTT2, A0QTT5, A0QTT6, A0QTW7, A0QTX2, A0QU01, A0QUE0, A0QUV5, A0QUX6, A0QUX7, A0QUY7, A0QV37, A0QV42, A0QV43, A0QV90, A0QVH8, A0QVH9, A0QVJ4, A0QVL1, A0QVL3, A0QVP3, A0QW36, A0QW41, A0QWH6, A0QWJ6, A0QWM9, A0QWQ6, A0QWR4, A0QWR5, A0QWS2, A0QWS3, A0QWS4, A0QWS5, A0QWU5, A0QWV6, A0QWX8, A0QX00, A0QX01, A0QX17, A0QX25, A0QX35, A0QX36, A0QX46, A0QX87, A0QXA2, A0QXA3, A0QXC0, A0QXP6, A0QXV0, A0QY23, A0QY48, A0QY89, A0QYD5, A0QYF1, A0QYH7, A0QYI2, A0QYS5, A0QYU6, A0QYX0, A0QZC9, A0QZR0, A0QZX4, A0R0C7, A0R0F7, A0R0H6, A0R0H7, A0R0I6, A0R0N6, A0R0S6, A0R0U5, A0R0V0, A0R112, A0R130, A0R148, A0R189, A0R198, A0R199, A0R1A9, A0R1X2, A0R203, A0R212, A0R214, A0R218, A0R293, A0R2B6, A0R2B7, A0R2J0, A0R2M2, A0R2N4, A0R2T3, A0R2T4, A0R2V7, A0R2W9, A0R2Y4, A0R318, A0R3F5, A0R3I8, A0R3L8, A0R3U1, A0R3W0, A0R436, A0R451, A0R479, A0R4B7, A0R4C3, A0R4C4, A0R4G8, A0R4I7, A0R4K5, A0R4Y7, A0R511, A0R5G4, A0R617, A0R635, A0R652, A0R665, A0R696, A0R699, A0R6E9, A0R6J9, A0R6M5, A0R6Z5, A0R710, A0R763, A0R7F4, A0R7F9, A0R7J0, P0CH36, P0CH37, P60281 |

| **Background (n = 3,181)** |
| --- |
| A0QND6, A0QND7, A0QND8, A0QND9, A0QNE0, A0QNE2, A0QNE6, A0QNE7, A0QNF1, A0QNF2, A0QNF5, A0QNF6, A0QNF7, A0QNF9, A0QNG0, A0QNG1, A0QNG2, A0QNG3, A0QNG4, A0QNG5, A0QNG6, A0QNG7, A0QNH2, A0QNH4, A0QNH5, A0QNH6, A0QNH8, A0QNH9, A0QNI0, A0QNI2, A0QNI3, A0QNI4, A0QNI7, A0QNI9, A0QNJ0, A0QNJ1, A0QNJ2, A0QNJ4, A0QNJ5, A0QNJ6, A0QNJ7, A0QNJ8, A0QNJ9, A0QNK0, A0QNK1, A0QNK2, A0QNK4, A0QNK6, A0QNK8, A0QNK9, A0QNL0, A0QNL1, A0QNL2, A0QNL3, A0QNL4, A0QNL5, A0QNL6, A0QNL9, A0QNM0, A0QNM1, A0QNM2, A0QNM4, A0QNM5, A0QNM6, A0QNM8, A0QNM9, A0QNN0, A0QNN6, A0QNN7, A0QZQ2, A0QNN8, A0QZQ3, A0QNN9, A0QNP1, A0QNP2, A0QNP3, A0QNP4, A0QNP6, A0QNP8, A0QNP9, A0QNQ4, A0QNQ5, A0QNQ6, A0QNQ7, A0QNQ9, A0QNR0, A0QNR3, A0QNR5, A0QNR6, A0QNR9, A0QNS1, A0QNS4, A0QNS6, A0QNT1, A0QNT2, A0QNT3, A0QNT6, A0QNU4, A0QNV7, A0QNW0, A0QNW6, A0QNX0, A0QNX2, A0QNY0, A0QNY3, A0QNY6, A0QNY8, A0QNY9, A0QNZ1, A0QNZ2, A0QNZ3, A0QNZ4, A0QNZ6, A0QNZ7, A0QNZ8, A0QNZ9, A0QP01, A0QP02, A0QP03, A0QP06, A0QP07, A0QP08, A0QP10, A0QP11, A0QP12, A0QP13, A0QP15, A0QP16, A0QP17, A0QP18, A0QP21, A0QP22, A0QP26, A0QP27, A0QP28, A0QP29, A0QP32, A0QP37, A0QP38, A0QP39, A0QP40, A0QP45, A0QP46, A0QP47, A0QP58, A0QP59, A0QP61, A0QP67, A0QP73, A0QP78, A0QP80, A0QP81, A0QP82, A0QP86, A0QP88, A0QP89, A0QP90, A0QP91, A0QP93, A0QP94, A0QP97, A0QPB3, A0QPC3, A0QPC7, A0QPD1, A0QPD4, A0QPD6, A0QPD7, A0QPD8, A0QPD9, A0QPE0, A0QPE1, A0QPE2, A0QPE3, A0QPE4, A0QPE5, A0QPE6, A0QPE7, A0QPE8, A0QPF6, A0QPF7, A0QPF9, A0QPG1, A0QPG2, A0QPG3, A0QPG4, A0QPG6, A0QPG7, A0QPG9, A0QPH0, A0QPH5, A0QPH6, A0QPI4, A0QPI9, A0QPJ0, A0QPJ1, A0QPJ3, A0QPJ7, A0QPL0, A0QPL1, A0QPL3, A0QPL6, A0QPL8, A0QPM0, A0QPM1, A0QPM2, A0QPM3, A0QPM7, A0QPM9, A0QPN0, A0QPN1, A0QPN2, A0QPN5, A0QPN9, A0QPP0, A0QPP1, A0QPP3, A0QPQ5, A0QPR6, A0QPS6, A0QPS9, A0QR29, A0R3I3, A0QPU4, A0QPV4, A0QPV5, A0QPV8, A0QPV9, A0QPW0, A0QPW1, A0QPW2, A0QPW3, A0QPW5, A0QPW6, A0QPW9, A0QPX0, A0QZL8, A0QPX1, A0R4N0, A0QW49, A0QPX2, A0QPX3, A0QPX4, A0QPX5, A0QPX6, A0QPY2, A0QPY4, A0QPY5, A0QPY6, A0QPY7, A0QQ02, A0QQ09, A0QQ15, A0QQ16, A0QQ17, A0QQ18, A0QQ19, A0QQ22, A0QQ23, A0QQ26, A0QQ27, A0QQ28, A0QQ30, A0QQ31, A0QQ36, A0QQ37, A0QQ38, A0QQ39, A0QQ40, A0QQ44, A0QQ46, A0QQ47, A0QQ48, A0QQ49, A0QQ51, A0QQ52, A0QQ54, A0QQ56, A0QQ58, A0QQ59, A0QQ60, A0QQ61, A0QQ62, A0QQ63, A0QQ64, A0QQ65, A0QQ67, A0QQ76, A0QQ77, A0QQ84, A0QQ90, A0QQ91, A0QQ93, A0QQ98, A0QQ99, A0QQA1, A0QQA2, A0QQA3, A0QQA8, A0QQB0, A0QQB1, A0QQB2, A0QQB4, A0QQB7, A0QQC1, A0QQC2, A0QQC7, A0QQC8, A0QQC9, A0QQD0, A0QQD2, A0QQD7, A0QQD8, A0QT56, A0QQE2, A0QQE9, A0QQF0, A0QQF1, A0QQF4, A0QQF5, A0QQF6, A0QQF7, A0QQF8, A0QQF9, A0QQG1, A0QQG9, A0QQH0, A0QQH1, A0QQH2, A0QQH5, A0QQH7, A0QQH8, A0QQI0, A0QQI1, A0QQI2, A0QQI3, A0QQI4, A0QQI5, A0QQI6, A0QQI7, A0QQI9, A0QQJ3, A0QQJ4, A0QQJ6, A0QQJ8, A0QQJ9, A0QQK0, A0QQK1, A0QQK3, A0QQK4, A0QQK5, A0QQK6, A0QQK7, A0QQK8, A0QQK9, A0QQL0, A0QQL1, A0QQL2, A0QQM2, A0QQM7, A0QQN0, A0QQN1, A0QQN5, A0QQN9, A0QQP0, A0QQP1, A0QQP2, A0QQP4, A0QQP5, A0QQP7, A0QQP8, A0QQP9, A0QQQ0, A0QQQ1, A0QQQ2, A0QQQ4, A0QQQ5, A0QQQ6, A0QQQ7, A0QQQ8, A0QQR0, A0QQR1, A0QQR2, A0QQR3, A0QQS3, A0QQS4, A0QQS5, A0QQS6, A0QQS7, A0QQS8, A0QQT1, A0QQT2, A0QQT9, A0QQU1, A0QQU2, A0QQU3, A0QQU5, A0QQU7, A0QQU8, A0QQV2, A0QQV4, A0QQV5, A0QQV7, A0QQV9, A0QQW0, A0QQW2, A0QQW3, A0QQW5, A0QQW6, A0QQW8, A0QQX0, A0QQX1, A0QQX4, A0QQX6, A0QQX7, A0QQX8, A0QQY0, A0QQY1, A0QQY2, A0QQY3, A0QQY7, A0QQY8, A0QQY9, A0QQZ1, A0QQZ2, A0QQZ3, A0QQZ4, A0QQZ5, A0QQZ8, A0QQZ9, A0QR00, A0QR01, A0QR03, A0QR04, A0QR05, A0QR06, A0QR07, A0QR08, A0QR09, A0QR11, A0QR12, A0QR13, A0QR14, A0QR17, A0QR18, A0QR19, A0QR20, A0QR24, A0QR26, A0QR28, A0QR31, A0QR33, A0QR34, A0QR35, A0QR37, A0QR38, A0QR39, A0QR40, A0QR41, A0QR42, A0QR46, A0QR48, A0QR49, A0QR50, A0QR51, A0QR53, A0QR54, A0QR55, A0QR56, A0QR65, A0QR68, A0QR69, A0QR72, A0QR73, A0QR74, A0QR76, A0QR77, A0QR78, A0QR79, A0QR80, A0QR82, A0QR87, A0QR88, A0QR89, A0QR90, A0QR91, A0QR94, A0QR98, A0QRA0, A0QRA2, A0QRA5, A0QRA6, A0QRA7, A0QRA8, A0QRA9, A0QRB0, A0QRB1, A0QRB3, A0QRB4, A0QRB6, A0QRB8, A0QRB9, A0QRC0, A0QRC1, A0QRC4, A0QRC5, A0QRC6, A0QRD1, A0QRD2, A0QRD3, A0QRD4, A0QRD5, A0QRD6, A0QRE7, A0QRF4, A0QRF5, A0QRF9, A0QRG0, A0QRG4, A0QRG5, A0QRG7, A0QRG8, A0QRH0, A0QRH1, A0QRH3, A0QRH9, A0QRI7, A0QRI8, A0QRI9, A0QRJ0, A0QRJ2, A0QRJ4, A0QRJ6, A0QRL0, A0QRL2, A0QRL3, A0QRL4, A0QRM0, A0QRN3, A0QRN4, A0QRN6, A0QRN7, A0QRP2, A0QRP3, A0QRP5, A0QRP8, A0QRP9, A0QRQ3, A0QRQ7, A0QRQ9, A0QRR0, A0QRR4, A0QRR5, A0QRR9, A0QRS0, A0QRS5, A0QRT0, A0QRT7, A0QRT9, A0QRU1, A0QRU5, A0QRU6, A0QRU7, A0QRU8, A0QRU9, A0QRV0, A0QRV1, A0QRV5, A0QRV6, A0QRV7, A0QRV8, A0QRW3, A0QRW4, A0QRW8, A0QRX0, A0QRX1, A0QRX9, A0QRY5, A0QRY6, A0QRY7, A0QRY8, A0QRZ0, A0QRZ3, A0QRZ8, A0QRZ9, A0QS07, A0QS13, A0QS18, A0QS19, A0QS24, A0QS28, A0QS29, A0QS30, A0QS33, A0QS34, A0QS36, A0QS38, A0QS39, A0QS40, A0QS41, A0QS42, A0QS43, A0QS44, A0QS45, A0QS46, A0QS49, A0QS51, A0QS52, A0QS54, A0QS61, A0QS62, A0QS63, A0QS64, A0QS66, A0QS67, A0QS73, A0QS74, A0QS77, A0QS78, A0QS79, A0QS80, A0QS81, A0QS85, A0QS86, A0QS88, A0QS90, A0QS91, A0QS92, A0QS95, A0QS96, A0QS97, A0QS98, A0QSA0, A0QSA5, A0QSA9, A0QSB0, A0QSB1, A0QSB2, A0QSB4, A0QSB7, A0QSB8, A0QSB9, A0QSC2, A0QSC4, A0QSC5, A0QSC8, A0QSD0, A0QSD1, A0QSD2, A0QSD3, A0QSD4, A0QSD5, A0QSD6, A0QSD7, A0QSD8, A0QSD9, A0QSE0, A0QSE2, A0QSE3, A0QSE5, A0QSF9, A0QSG0, A0QSG1, A0QSG2, A0QSG3, A0QSG4, A0QSG5, A0QSG6, A0QSG7, A0QSG8, A0QSG9, A0QSH0, A0QSH1, A0QSH3, A0QSH4, A0QSH5, A0QSH6, A0QSH7, A0QSH8, A0QSH9, A0QSI0, A0QSI6, A0QSI7, A0QSI8, A0QSJ0, A0QSJ1, A0QSJ2, A0QSK5, A0QSK6, A0QSK7, A0QSK8, A0QSK9, A0QSL0, A0QSL1, A0QSL3, A0QSL5, A0QSL6, A0QSL7, A0QSL8, A0QSL9, A0QSM0, A0QSM1, A0QSM2, A0QSM3, A0QSM5, A0QSN4, A0QSN6, A0QSN7, A0QSN8, A0QSN9, A0QSP0, A0QSP1, A0QSP2, A0QSP5, A0QSP6, A0QSP7, A0QSP8, A0QSP9, A0QSQ1, A0QSQ4, A0QSQ6, A0QSQ7, A0QSQ8, A0QSR0, A0QSR4, A0QSR5, A0QSR6, A0QSR7, A0QSR8, A0QSR9, A0QSS0, A0QSS1, A0QSS2, A0QSS3, A0QSS4, A0QST4, A0QST9, A0QSU0, A0QSU3, A0QSU4, A0QSU5, A0QSV0, A0QSV1, A0QSW1, A0QSX2, A0QSX3, A0QSX4, A0QSX6, A0QSX7, A0QSX9, A0QSY0, A0QSY1, A0QSY2, A0QSY4, A0QSY5, A0QSY9, A0QSZ0, A0QSZ1, A0QSZ3, A0QSZ4, A0QSZ5, A0QSZ6, A0QSZ7, A0QSZ9, A0QT00, A0QT01, A0QT02, A0QT04, A0QT05, A0QT07, A0QT08, A0QT09, A0QT10, A0QT11, A0QT13, A0QT14, A0QT16, A0QT17, A0QT18, A0QT19, A0QT20, A0QT21, A0QT22, A0QT32, A0QT33, A0QT39, A0QT40, |

A0QT41, A0QT45, A0QT46, A0QT58, A0QT68, A0QT69, A0QT70, A0QT72, A0QT73, A0QT74, A0QT75, A0QT76, A0QT77, A0QT79, A0QT80, A0QT84, A0QT87, A0QT91, A0QT95, A0QT96, A0QT97, A0QT98, A0QTA4, A0QTA5, A0QTA7, A0QTB1, A0QTC7, A0QTD7, A0QTD8, A0QTE0, A0QTE1, A0QTE2, A0QTE3, A0QTE5, A0QTE6, A0QTE7, A0QTF0, A0QTF1, A0QTF2, A0QTF3, A0QTF4, A0QTF5, A0QTF6, A0QTF7, A0QTF8, A0QTF9, A0QTG1, A0QTG2, A0QTG3, A0QTG4, A0QTG6, A0QTG7, A0QTG8, A0QTG9, A0QTH0, A0QTH4, A0QTH9, A0QTI0, A0QTI1, A0QTI2, A0QTI3, A0QTI9, A0QTJ7, A0QTK1, A0QTK2, A0QTK3, A0QTK4, A0QTK6, A0QTL0, A0QTL1, A0QTL2, A0QTL3, A0QTL4, A0QTL6, A0QTL7, A0QTL8, A0QTL9, A0QTM0, A0QTM1, A0QTM2, A0QTM9, A0QTN0, A0QTP1, A0QTP2, A0QTP6, A0QTP7, A0QTP8, A0QTQ0, A0QTQ2, A0QTQ3, A0QTQ4, A0QTQ5, A0QTQ6, A0QTQ7, A0QTQ8, A0QTQ9, A0QTR1, A0QTR2, A0QTR3, A0QTR4, A0QTR5, A0QTR6, A0QTR7, A0QTR8, A0QTS0, A0QTS1, A0QTS2, A0QTS3, A0QTS4, A0QTS5, A0QTS8, A0QTS9, A0QTT0, A0QTT2, A0QTT5, A0QTT6, A0QTT7, A0QTU1, A0QTU3, A0QTV3, A0QTV6, A0QTV7, A0QTV8, A0QTW2, A0QTW7, A0QTX1, A0QTX4, A0QTY2, A0QTY3, A0QTY5, A0QTY6, A0QTZ0, A0QTZ1, A0QTZ2, A0QTZ3, A0QTZ4, A0QTZ9, A0QU00, A0QU01, A0QU03, A0QU06, A0QU07, A0QU10, A0QU11, A0QU12, A0QU15, A0QU17, A0QU18, A0QU19, A0QU20, A0QU21, A0QU34, A0QU37, A0QU39, A0QU42, A0QU43, A0QU45, A0QU46, A0QU47, A0QU48, A0QU51, A0QU52, A0QU53, A0QU54, A0QU56, A0QU57, A0QU58, A0QU59, A0QU61, A0QU62, A0QU63, A0QU64, A0QU69, A0QU77, A0QU80, A0QU81, A0QU82, A0QU86, A0QU87, A0QU88, A0QU89, A0QU91, A0QU92, A0QU93, A0QU95, A0QU97, A0QU98, A0QUA0, A0QUA1, A0QUA2, A0QUA6, A0QUC0, A0QUC9, A0QUD0, A0QUD9, A0QUE0, A0QUE1, A0QUE5, A0QUF4, A0QUF8, A0QUF9, A0QUG0, A0QUG1, A0QUG2, A0QUG3, A0QUG4, A0QUG6, A0QUG7, A0QUH0, A0QUH1, A0QUH2, A0QUH3, A0QUH4, A0QUH6, A0QUH8, A0QUI8, A0QUI9, A0QUJ1, A0QUK7, A0QUL0, A0QUL2, A0QUL6, A0QUM3, A0QUM5, A0QUM6, A0QUM7, A0QUM8, A0QUN1, A0QUN2, A0QUN3, A0QUN4, A0QUN5, A0QUN7, A0QUN8, A0QUN9, A0QUP0, A0QUP3, A0QUP4, A0QZB5, A0QUV4, A0QUV5, A0QUV6, A0QUV7, A0QUV8, A0QUW2, A0QUW3, A0QUW4, A0QUW5, A0QUW6, A0QUW7, A0QUW8, A0QUX0, A0QUX1, A0QUX3, A0QUX4, A0QUX5, A0QUX6, A0QUX7, A0QUX8, A0QUX9, A0QUY0, A0QUY1, A0QUY2, A0QUY3, A0QUY5, A0QUY6, A0QUY7, A0QUY8, A0QUY9, A0QUZ0, A0QUZ2, A0QUZ3, A0QUZ4, A0QUZ5, A0QUZ7, A0QUZ8, A0QUZ9, A0QV00, A0QV01, A0QV03, A0QV04, A0QV05, A0QV09, A0QV10, A0QV11, A0QV12, A0QV14, A0QV15, A0QV16, A0QV17, A0QV18, A0QV19, A0QV20, A0QV21, A0QV23, A0QV24, A0QV25, A0QV26, A0QV28, A0QV29, A0QV31, A0QV32, A0QV33, A0QV35, A0QV36, A0QV37, A0QV38, A0QV39, A0QV40, A0QV41, A0QV42, A0QV43, A0QV45, A0QV46, A0QV47, A0QV49, A0QV51, A0QV52, A0QV53, A0QV55, A0QV56, A0QV58, A0QV60, A0QV61, A0QV67, A0QV69, A0QV70, A0QV71, A0QV74, A0QV76, A0QV77, A0QV88, A0QV89, A0QV90, A0QV91, A0QV92, A0QV97, A0QV98, A0QVB1, A0QVB2, A0QVB5, A0QVB6, A0QVB8, A0QVB9, A0QVC0, A0QVC7, A0QVC8, A0QVC9, A0QVD1, A0QVD4, A0QVD5, A0QVD7, A0QVD8, A0QVD9, A0QVE0, A0QVE4, A0QVE5, A0QVF2, A0QVF3, A0QVF6, A0QVH0, A0QVH7, A0QVH8, A0QVH9, A0QVI1, A0QVI3, A0QVI4, A0QVI6, A0QVI7, A0QVI8, A0QVJ0, A0QVJ2, A0QVJ3, A0QVJ4, A0QVJ5, A0QVJ6, A0QVJ7, A0QVK4, A0QVK5, A0QVL0, A0QVL1, A0QVL2, A0QVL3, A0QVL4, A0QVL5, A0QVL6, A0QVL7, A0QVL8, A0QVL9, A0QVM0, A0QVM1, A0QVM2, A0QVM3, A0QVM4, A0QVM5, A0QVM7, A0QVM8, A0QVM9, A0QVP0, A0QVP1, A0QVP2, A0QVP3, A0QVP4, A0QVP6, A0QVP7, A0QVP8, A0QVP9, A0QVQ0, A0QVQ1, A0QVQ2, A0QVQ3, A0QVQ5, A0QVQ6, A0QVQ7, A0QVQ8, A0QVQ9, A0QVR0, A0QVR3, A0QVR4, A0QVR5, A0QVR6, A0QVR7, A0QVR8, A0QVR9, A0QVS0, A0QVS1, A0QVS6, A0QVS9, A0QVT0, A0QVT1, A0QVT2, A0QVT4, A0QVT5, A0QVT6, A0QVT8, A0QVT9, A0QVU0, A0QVU2, A0QVU3, A0QVU4, A0QVU5, A0QVU6, A0QVU7, A0QVX1, A0QVX2, A0QVX3, A0QVX4, A0QVX5, A0QVX6, A0QVX8, A0QVX9, A0QVY0, A0QVY1, A0QVY3, A0QVY4, A0QVY5, A0QVY8, A0QVY9, A0QVZ0, A0QVZ3, A0QVZ5, A0QVZ6, A0QVZ9, A0QW02, A0QW03, A0QW04, A0QW05, A0QW06, A0QW07, A0QW08, A0QW09, A0QW10, A0QW11, A0QW12, A0QW13, A0QW14, A0QW15, A0QW16, A0QW17, A0QW19, A0QW20, A0QW21, A0QW22, A0QW23, A0QW24, A0QW25, A0QW27, A0QW28, A0QW29, A0QW30, A0QW31, A0QW32, A0QW34, A0QW35, A0QW36, A0QW37, A0QW41, A0QW43, A0QW47, A0QW54, A0QW60, A0QW62, A0QW71, A0QW82, A0QWA0, A0QWB3, A0QWB5, A0QWC7, A0QWD0, A0QWD2, A0QWD3, A0QWD4, A0QWE1, A0QWE4, A0QWE9, A0QWF0, A0QWF1, A0QWF4, A0QWF7, A0QWF9, A0QWG0, A0QWG2, A0QWG3, A0QWG4, A0QWG5, A0QWG6, A0QWG7, A0QWG8, A0QWG9, A0QWH0, A0QWH1, A0QWH2, A0QWH3, A0QWH4, A0QWH5, A0QWH6, A0QWI4, A0QWI5, A0QWI7, A0QWI8, A0QWJ1, A0QWJ2, A0QWJ3, A0QWJ4, A0QWJ5, A0QWJ6, A0QWK2, A0QWK4, A0QWK5, A0QWK6, A0QWK7, A0QWL3, A0QWL4, A0QWL9, A0QWM8, A0QWM9, A0QWN0, A0QWN1, A0QWN3, A0QWN4, A0QWP3, A0QWP6, A0QWP7, A0QWP8, A0QWP9, A0QWQ0, A0QWQ1, A0QWQ3, A0QWQ4, A0QWQ5, A0QWQ6, A0QWQ7, A0QWQ9, A0QWR0, A0QWR1, A0QWR3, A0QWR4, A0QWR5, A0QWR8, A0QWR9, A0QWS0, A0QWS1, A0QWS2, A0QWS3, A0QWS4, A0QWS5, A0QWS8, A0QWS9, A0QWT1, A0QWT2, A0QWT3, A0QWT4, A0QWT5, A0QWT6, A0QWT7, A0QWT9, A0QWU0, A0QWU1, A0QWU2, A0QWU3, A0QWU4, A0QWU5, A0QWU7, A0QWU8, A0QWU9, A0QWV0, A0QWV1, A0QWV2, A0QWV4, A0QWV6, A0QWV7, A0QWV8, A0QWV9, A0QWW2, A0QWW3, A0QWW4, A0QWW5, A0QWX1, A0QWX3, A0QWX4, A0QWX6, A0QWX7, A0QWX8, A0QWX9, A0QWY0, A0QWY2, A0QWY3, A0QWY4, A0QWY8, A0QWY9, A0QWZ0, A0QWZ3, A0QWZ4, A0QWZ6, A0QWZ7, A0QWZ8, A0QWZ9, A0QX00, A0QX01, A0QX02, A0QX03, A0QX04, A0QX14, A0QX15, A0QX16, A0QX17, A0QX19, A0QX20, A0QX21, A0QX24, A0QX25, A0QX26, A0QX29, A0QX30, A0QX31, A0QX32, A0QX33, A0QX35, A0QX36, A0QX37, A0QX45, A0QX46, A0QX47, A0QX48, A0QX50, A0QX51, A0QX52, A0QX55, A0QX60, A0QX61, A0QX62, A0QX63, A0QX64, A0QX65, A0QX66, A0QX69, A0QX70, A0QX72, A0QX74, A0QX75, A0QX76, A0QX77, A0QX80, A0QX81, A0QX82, A0QX83, A0QX84, A0QX85, A0QX86, A0QX87, A0QX88, A0QX90, A0QX91, A0QX92, A0QX93, A0QX94, A0QX95, A0QX96, A0QX97, A0QX98, A0QXA1, A0QXA2, A0QXA3, A0QXA4, A0QXA5, A0QXA6, A0QXA7, A0QXA8, A0QXA9, A0QXB0, A0QXB1, A0QXB2, A0QXB3, A0QXB4, A0QXB5, A0QXB9, A0QXC0, A0QXC1, A0QXC2, A0QXC3, A0QXC4, A0QXC6, A0QXC7, A0QXC8, A0QXD0, A0QXD2, A0QXD4, A0QXD5, A0QXD6, A0QXD7, A0QXD8, A0QXD9, A0QXE0, A0QXE2, A0QXE3, A0QXE4, A0QXE5, A0QXF0, A0QXF1, A0QXF3, A0QXF4, A0QXF7, A0QXF8, A0QXF9, A0QXG0, A0QXH0, A0QXH3, A0QXH4, A0QXH8, A0QXH9, A0QXI0, A0QXI4, A0QXI7, A0QXI9, A0QXJ0, A0QXJ8, A0QXK2, A0QXK4, A0QXK5, A0QXL4, A0QXL5, A0QXL6, A0QXM5, A0QXM6, A0QXM7, A0QXM8, A0QXM9, A0QXN2, A0QXP3, A0QXP4, A0QXP6, A0QXP8, A0QXP9, A0QXQ2, A0QXR2, A0QXR3, A0QXR4, A0QXS3, A0QXS8, A0QXT4, A0QXT5, A0QXT8, A0QXU0, A0QXU2, A0QXU6, A0QXV0, A0QXV1, A0QXV6, A0QXV8, A0QXW6, A0QXW7, A0QXX1, A0QXX3,

A0QXX5, A0QXX7, A0QXY1, A0QXY3, A0QXY5, A0QXY6, A0QXY7, A0QXZ2, A0QXZ3, A0QXZ4, A0QXZ5, A0QXZ6, A0QXZ9, A0QY04, A0QY05, A0QY08, A0QY09, A0QY11, A0QY12, A0QY16, A0QY21, A0QY22, A0QY23, A0QY24, A0QY28, A0QY29, A0QY30, A0QY31, A0QY32, A0QY34, A0QY36, A0QY40, A0QY41, A0QY42, A0QY44, A0QY49, A0QY50, A0QY53, A0QY56, A0QY73, A0QY76, A0QY79, A0QY81, A0QY83, A0QY84, A0QY85, A0QY89, A0QY90, A0QY91, A0QY95, A0QY96, A0QYA7, A0QYA9, A0QYB0, A0QYB1, A0QYB3, A0QYB5, A0QYB7, A0QYB9, A0QYC0, A0QYC1, A0QYC2, A0QYC3, A0QYC5, A0QYC8, A0QYD3, A0QYD4, A0QYD5, A0QYD6, A0QYD7, A0QYD8, A0QYD9, A0QYE0, A0QYE1, A0QYE2, A0QYE3, A0QYE5, A0QYE7, A0QYE8, A0QYF1, A0QYF2, A0QYF3, A0QYF4, A0QYF5, A0QYF6, A0QYF7, A0QYF9, A0QYG0, A0QYG1, A0QYG2, A0QYG3, A0QYG4, A0QYG6, A0QYG9, A0QYH0, A0QYH1, A0QYH2, A0QYH4, A0QYH5, A0QYH6, A0QYH7, A0QYH8, A0QYI1, A0QYI2, A0QYI8, A0QYJ2, A0QYJ3, A0QYJ5, A0QYJ6, A0QYJ8, A0QYJ9, A0QYK0, A0QYK2, A0QYK4, A0QYK5, A0QYK6, A0QYK8, A0QYL7, A0QYL8, A0QYL9, A0QYM1, A0QYP5, A0QYP9, A0QYQ0, A0QYQ1, A0QYQ2, A0QYQ3, A0QYQ4, A0QYQ5, A0QYQ6, A0QYQ7, A0QYQ8, A0QYQ9, A0QYR0, A0QYR2, A0QYR3, A0QYR4, A0QYR5, A0QYS0, A0QYS1, A0QYS2, A0QYS3, A0QYS4, A0QYS6, A0QYS7, A0QYS8, A0QYS9, A0QYT0, A0QYT1, A0QYT2, A0QYT3, A0QYT4, A0QYT5, A0QYT7, A0QYT9, A0QYU0, A0QYU1, A0QYU2, A0QYU3, A0QYU4, A0QYU5, A0QYU6, A0QYU8, A0QYV0, A0QYV1, A0QYV3, A0QYW0, A0QYW2, A0QYW4, A0QYW5, A0QYW6, A0QYW7, A0QYX0, A0QYX2, A0QYX6, A0QYY3, A0QYY4, A0QYY6, A0QYZ0, A0QYZ1, A0QYZ2, A0QYZ6, A0QYZ7, A0QZ01, A0QZ03, A0QZ04, A0QZ05, A0QZ08, A0QZ09, A0QZ11, A0QZ12, A0QZ13, A0QZ14, A0QZ16, A0QZ17, A0QZ24, A0QZ25, A0QZ26, A0QZ29, A0QZ30, A0QZ31, A0QZ32, A0QZ33, A0QZ34, A0QZ35, A0QZ36, A0QZ37, A0QZ38, A0QZ39, A0QZ40, A0QZ41, A0QZ42, A0QZ47, A0QZ48, A0QZ49, A0QZ50, A0QZ51, A0QZ52, A0QZ54, A0QZ55, A0QZ56, A0QZ57, A0QZ58, A0QZ59, A0QZ60, A0QZ61, A0QZ62, A0QZ65, A0QZ66, A0QZ69, A0QZ78, A0QZ79, A0QZ82, A0QZ83, A0QZ84, A0QZ85, A0QZ86, A0QZ91, A0QZ92, A0QZ93, A0QZ95, A0QZ96, A0QZ97, A0QZ98, A0QZ99, A0QZA1, A0QZA2, A0QZA6, A0QZA9, A0QZB0, A0QZB2, A0QZB3, A0QZC6, A0QZC7, A0QZC9, A0QZD1, A0QZD6, A0QZF0, A0QZF5, A0QZH0, A0QZH7, A0QZH8, A0QZI2, A0QZJ0, A0QZJ1, A0QZJ2, A0QZJ5, A0QZK5, A0QZK7, A0QZL1, A0QZM4, A0QZM6, A0QZM7, A0QZM8, A0QZM9, A0QZN0, A0QZN1, A0QZN7, A0QZP1, A0QZQ4, A0QZQ6, A0QZQ7, A0QZQ8, A0QZQ9, A0QZR0, A0QZR1, A0QZR2, A0QZR4, A0QZR5, A0QZR7, A0QZR9, A0QZS0, A0QZS2, A0QZT3, A0QZT4, A0QZT7, A0QZU8, A0QZV4, A0QZV7, A0QZW4, A0QZW7, A0QZW8, A0QZX1, A0QZX2, A0QZX4, A0QZX6, A0QZX7, A0QZX8, A0QZX9, A0QZY0, A0QZY1, A0QZY2, A0QZY3, A0QZY4, A0QZY6, A0QZY7, A0QZY9, A0QZZ0, A0QZZ1, A0R003, A0R005, A0R006, A0R008, A0R009, A0R010, A0R012, A0R014, A0R015, A0R016, A0R018, A0R019, A0R020, A0R021, A0R024, A0R025, A0R026, A0R027, A0R028, A0R029, A0R030, A0R031, A0R032, A0R033, A0R034, A0R035, A0R036, A0R037, A0R039, A0R040, A0R041, A0R042, A0R043, A0R044, A0R045, A0R048, A0R049, A0R050, A0R051, A0R052, A0R053, A0R054, A0R057, A0R058, A0R059, A0R060, A0R061, A0R062, A0R063, A0R064, A0R066, A0R067, A0R069, A0R070, A0R071, A0R072, A0R073, A0R074, A0R075, A0R076, A0R078, A0R079, A0R080, A0R082, A0R083, A0R084, A0R085, A0R087, A0R088, A0R089, A0R090, A0R092, A0R093, A0R094, A0R095, A0R096, A0R097, A0R098, A0R099, A0R0A0, A0R0A1, A0R0A4, A0R0A7, A0R0A8, A0R0A9, A0R0B0, A0R0B1, A0R0B2, A0R0B3, A0R0B4, A0R0B5, A0R0B6, A0R0B7, A0R0C0, A0R0C1, A0R0C3, A0R0C4, A0R0C8, A0R0D0, A0R0D4, A0R0D5, A0R0D6, A0R0D8, A0R0E0, A0R0E5, A0R0E7, A0R0E9, A0R0F3, A0R0F4, A0R0F7, A0R0G8, A0R0H2, A0R0H6, A0R0H7, A0R0H9, A0R0I1, A0R0I3, A0R0I6, A0R0I7, A0R0I8, A0R0M4, A0R0N5, A0R0N6, A0R0N9, A0R0P8, A0R0P9, A0R0Q4, A0R0Q5, A0R0Q6, A0R0Q9, A0R0R0, A0R0R1, A0R0R3, A0R0R8, A0R0R9, A0R0S0, A0R0S1, A0R0S2, A0R0S3, A0R0S4, A0R0S5, A0R0S6, A0R0S7, A0R0S8, A0R0S9, A0R0T0, A0R0T1, A0R0T7, A0R0T8, A0R0T9, A0R0U1, A0R0U3, A0R0U4, A0R0U5, A0R0U6, A0R0U7, A0R0U8, A0R0U9, A0R0V0, A0R0V2, A0R0V8, A0R0V9, A0R0W1, A0R0W2, A0R0W4, A0R0W5, A0R0W6, A0R0W7, A0R0W9, A0R0X1, A0R0X2, A0R0X9, A0R0Y6, A0R0Y9, A0R0Z0, A0R0Z1, A0R0Z2, A0R0Z3, A0R0Z4, A0R0Z5, A0R0Z8, A0R0Z9, A0R100, A0R101, A0R102, A0R103, A0R108, A0R109, A0R110, A0R111, A0R112, A0R113, A0R114, A0R115, A0R116, A0R117, A0R120, A0R125, A0R129, A0R130, A0R143, A0R145, A0R147, A0R148, A0R149, A0R150, A0R151, A0R152, A0R155, A0R156, A0R157, A0R158, A0R165, A0R169, A0R170, A0R171, A0R172, A0R175, A0R177, A0R178, A0R181, A0R183, A0R184, A0R185, A0R187, A0R189, A0R191, A0R192, A0R193, A0R194, A0R196, A0R197, A0R198, A0R199, A0R1A2, A0R1A4, A0R1A5, A0R1A6, A0R1A7, A0R1A8, A0R1A9, A0R1B0, A0R1B1, A0R1B3, A0R1B5, A0R1B6, A0R1B8, A0R1B9, A0R1C0, A0R1C1, A0R1C2, A0R1C3, A0R1C5, A0R1C6, A0R1C7, A0R1C8, A0R1C9, A0R1D0, A0R1D1, A0R1D5, A0R1D6, A0R1D7, A0R1D8, A0R1D9, A0R1E0, A0R1E1, A0R1E4, A0R1E5, A0R1E6, A0R1F2, A0R1G0, A0R1G3, A0R1G6, A0R1H2, A0R1H3, A0R1H5, A0R1H6, A0R1H7, A0R1I5, A0R1I9, A0R1J0, A0R1J2, A0R1J8, A0R1P4, A0R1P5, A0R1P6, A0R1P7, A0R1Q0, A0R1Q1, A0R1Q2, A0R1R5, A0R1R7, A0R1S4, A0R1V9, A0R1W1, A0R1W4, A0R1W5, A0R1W6, A0R1W7, A0R1W8, A0R1W9, A0R1X0, A0R1X2, A0R1X3, A0R1X4, A0R1X5, A0R1X6, A0R1X7, A0R1X8, A0R1X9, A0R1Y0, A0R1Y1, A0R1Y2, A0R1Y4, A0R1Y5, A0R1Y6, A0R1Y7, A0R1Y8, A0R1Z0, A0R1Z1, A0R1Z6, A0R1Z7, A0R1Z8, A0R1Z9, A0R200, A0R201, A0R202, A0R203, A0R204, A0R206, A0R207, A0R212, A0R213, A0R214, A0R215, A0R217, A0R218, A0R219, A0R220, A0R221, A0R226, A0R229, A0R230, A0R234, A0R237, A0R238, A0R239, A0R240, A0R241, A0R242, A0R248, A0R249, A0R253, A0R258, A0R260, A0R261, A0R266, A0R267, A0R268, A0R269, A0R271, A0R273, A0R277, A0R278, A0R280, A0R281, A0R283, A0R284, A0R287, A0R290, A0R291, A0R292, A0R293, A0R295, A0R298, A0R2A4, A0R2A5, A0R2A6, A0R2A8, A0R2B0, A0R2B1, A0R2B2, A0R2B3, A0R2B5, A0R2B6, A0R2B7, A0R2B8, A0R2B9, A0R2C0, A0R2C1, A0R2C2, A0R2C3, A0R2C4, A0R2C6, A0R2C7, A0R2C8, A0R2D0, A0R2D1, A0R2D2, A0R2D3, A0R2D4, A0R2D5, A0R2D6, A0R2E1, A0R2E2, A0R2E3, A0R2E4, A0R2E5, A0R2E6, A0R2E7, A0R2E8, A0R2E9, A0R2F0, A0R2F1, A0R2F2, A0R2F3, A0R2G2, A0R2G4, A0R2G5, A0R2H4, A0R2H5, A0R2H6, A0R2H8, A0R2H9, A0R2I0, A0R2I2, A0R2I3, A0R2I4, A0R2I7, A0R2I8, A0R2J0, A0R2J4, A0R2K3, A0R2K6, A0R2K7, A0R2L3, A0R2L4, A0R2L7, A0R2M2, A0R2M8, A0R2N0, A0R2N1, A0R2N3, A0R2N4, A0R2N5, A0R2P0, A0R2P1, A0R2P2, A0R2P3, A0R2P7, A0R2Q0, A0R2Q4, A0R2Q5, A0R2Q6, A0R2Q7, A0R2Q8, A0R2R4, A0R2R5, A0R2R6, A0R2R7, A0R2R8, A0R2R9, A0R2S1, A0R2S3, A0R2S4, A0R2S8, A0R2S9, A0R2T0, A0R2T1, A0R2T2, A0R2T3, A0R2T4, A0R2T6, A0R2U2, A0R2U5, A0R2U6, A0R2U7, A0R2U8, A0R2U9, A0R2V0, A0R2V1, A0R2V2, A0R2V3, A0R2V4, A0R2V5, A0R2V6, A0R2V7, A0R2V8, A0R2V9, A0R2W3, A0R2W4, A0R2W5, A0R2W6, A0R2W7, A0R2W9, A0R2X0, A0R2X1, A0R2X3, A0R2X4, A0R2X6, A0R2X8, A0R2Y0, A0R2Y1, A0R2Y2, A0R2Y3, A0R2Y4, A0R2Y5, A0R2Y6, A0R2Y7, A0R2Y8, A0R2Z0, A0R2Z1, A0R2Z2, A0R2Z3, A0R2Z4, A0R2Z5, A0R2Z9, A0R308, A0R310, A0R315, A0R316, A0R318,

A0R325, A0R326, A0R327, A0R336, A0R338, A0R344, A0R346, A0R351, A0R355, A0R358, A0R365, A0R368, A0R369, A0R373, A0R374, A0R376, A0R379, A0R380, A0R381, A0R391, A0R3A3, A0R3A4, A0R3A6, A0R3A8, A0R3B3, A0R3B5, A0R3B6, A0R3B7, A0R3B8, A0R3B9, A0R3C1, A0R3C2, A0R3C4, A0R3C5, A0R3C6, A0R3C7, A0R3C8, A0R3C9, A0R3D0, A0R3D1, A0R3D2, A0R3D3, A0R3D6, A0R3D7, A0R3D9, A0R3E1, A0R3E2, A0R3E3, A0R3E4, A0R3E5, A0R3E7, A0R3E8, A0R3F0, A0R3F1, A0R3F3, A0R3F5, A0R3F8, A0R3G9, A0R3H0, A0R3H1, A0R3H5, A0R3H7, A0R3H9, A0R3I2, A0R3I5, A0R3I6, A0R3I7, A0R3I8, A0R3I9, A0R3J4, A0R3K4, A0R3K5, A0R3L0, A0R3L1, A0R3L2, A0R3L3, A0R3L4, A0R3L5, A0R3L6, A0R3L8, A0R3L9, A0R3M0, A0R3M2, A0R3M3, A0R3M4, A0R3M6, A0R3N3, A0R3N5, A0R3N8, A0R3N9, A0R3P0, A0R3P3, A0R3P4, A0R3P7, A0R3Q0, A0R3Q1, A0R3Q2, A0R3Q3, A0R3R3, A0R3R5, A0R3R8, A0R3S0, A0R3S1, A0R3S3, A0R3S7, A0R3T1, A0R3T6, A0R3T7, A0R3T8, A0R3T9, A0R3U0, A0R3U1, A0R3V8, A0R3W0, A0R3X5, A0R3Y0, A0R3Y1, A0R3Y2, A0R3Y3, A0R3Y4, A0R3Y5, A0R3Y8, A0R3Z0, A0R3Z1, A0R401, A0R402, A0R403, A0R404, A0R405, A0R407, A0R408, A0R409, A0R411, A0R412, A0R414, A0R416, A0R417, A0R418, A0R420, A0R421, A0R422, A0R423, A0R425, A0R426, A0R427, A0R429, A0R430, A0R432, A0R433, A0R435, A0R436, A0R438, A0R439, A0R440, A0R441, A0R442, A0R443, A0R444, A0R446, A0R447, A0R448, A0R449, A0R450, A0R451, A0R452, A0R453, A0R458, A0R461, A0R462, A0R464, A0R465, A0R466, A0R469, A0R470, A0R471, A0R472, A0R473, A0R477, A0R478, A0R479, A0R480, A0R481, A0R484, A0R498, A0R4A8, A0R4A9, A0R4B3, A0R4B5, A0R4B6, A0R4B7, A0R4B8, A0R4C0, A0R4C1, A0R4C2, A0R4C3, A0R4C4, A0R4C5, A0R4C6, A0R4C8, A0R4C9, A0R4D0, A0R4D2, A0R4D5, A0R4D6, A0R4D7, A0R4D8, A0R4E0, A0R4F6, A0R4F7, A0R4G2, A0R4G3, A0R4G4, A0R4G7, A0R4G8, A0R4G9, A0R4H0, A0R4H1, A0R4H2, A0R4H3, A0R4H4, A0R4H6, A0R4H7, A0R4H9, A0R4I0, A0R4I3, A0R4I5, A0R4I6, A0R4I7, A0R4J1, A0R4J3, A0R4J7, A0R4J8, A0R4K1, A0R4K2, A0R4K4, A0R4K5, A0R4K9, A0R4L0, A0R4L1, A0R4L2, A0R4L6, A0R4L9, A0R4M2, A0R4M3, A0R4M4, A0R4M7, A0R4M8, A0R4M9, A0R4N1, A0R4N2, A0R4N3, A0R4N4, A0R4N5, A0R4N6, A0R4N7, A0R4N8, A0R4N9, A0R4P0, A0R4P1, A0R4P4, A0R4P5, A0R4P7, A0R4Q0, A0R4Q1, A0R4Q2, A0R4Q3, A0R4Q8, A0R4Q9, A0R4R0, A0R4R1, A0R4R2, A0R4R3, A0R4R9, A0R4S0, A0R4S3, A0R4S6, A0R4S7, A0R4S8, A0R4S9, A0R4T0, A0R4U4, A0R4V6, A0R4X1, A0R4X6, A0R4X7, A0R4Y1, A0R4Y2, A0R4Y4, A0R4Y7, A0R4Y9, A0R4Z0, A0R4Z1, A0R4Z2, A0R4Z5, A0R4Z6, A0R4Z8, A0R4Z9, A0R501, A0R502, A0R503, A0R507, A0R511, A0R514, A0R515, A0R517, A0R518, A0R522, A0R523, A0R524, A0R525, A0R527, A0R528, A0R529, A0R530, A0R532, A0R533, A0R535, A0R538, A0R539, A0R543, A0R546, A0R548, A0R555, A0R557, A0R558, A0R559, A0R560, A0R561, A0R562, A0R564, A0R565, A0R566, A0R568, A0R569, A0R570, A0R572, A0R573, A0R574, A0R576, A0R577, A0R579, A0R580, A0R581, A0R582, A0R584, A0R585, A0R586, A0R587, A0R588, A0R589, A0R590, A0R592, A0R593, A0R595, A0R596, A0R597, A0R5A8, A0R5A9, A0R5B0, A0R5B1, A0R5B2, A0R5B3, A0R5B4, A0R5B7, A0R5B8, A0R5C0, A0R5C1, A0R5C3, A0R5C4, A0R5C5, A0R5D3, A0R5D5, A0R5D6, A0R5D9, A0R5E0, A0R5E1, A0R5E2, A0R5E5, A0R5F5, A0R5F9, A0R5G1, A0R5G2, A0R5G4, A0R5G5, A0R5G6, A0R5G7, A0R5G8, A0R5G9, A0R5H1, A0R5H2, A0R5H3, A0R5H4, A0R5H5, A0R5H6, A0R5I0, A0R5I3, A0R5I4, A0R5I5, A0R5I8, A0R5I9, A0R5J3, A0R5J4, A0R5J5, A0R5J6, A0R5J8, A0R5K0, A0R5K1, A0R5K2, A0R5K4, A0R5K8, A0R5K9, A0R5L0, A0R5L3, A0R5L5, A0R5L6, A0R5L8, A0R5M1, A0R5M2, A0R5M3, A0R5M4, A0R5M7, A0R5M8, A0R5M9, A0R5N0, A0R5N4, A0R5N7, A0R5N8, A0R5N9, A0R5P4, A0R5P6, A0R5Q0, A0R5Q2, A0R5Q3, A0R5Q4, A0R5Q6, A0R5Q7, A0R5Q8, A0R5Q9, A0R5R0, A0R5R1, A0R5R2, A0R5R3, A0R5R4, A0R5R5, A0R5R6, A0R5R7, A0R5R8, A0R5R9, A0R5S0, A0R5S1, A0R5S3, A0R5S7, A0R5T0, A0R5T1, A0R5T2, A0R5T4, A0R5T6, A0R5T7, A0R5T8, A0R5U1, A0R5U2, A0R5U3, A0R5U4, A0R5U5, A0R5U7, A0R5U8, A0R5V1, A0R5V7, A0R5V8, A0R5V9, A0R5W5, A0R5W6, A0R5W7, A0R5W8, A0R5W9, A0R5X3, A0R5X8, A0R5X9, A0R5Y0, A0R5Y1, A0R5Y8, A0R5Y9, A0R5Z0, A0R5Z1, A0R5Z2, A0R5Z3, A0R5Z8, A0R5Z9, A0R606, A0R607, A0R609, A0R610, A0R611, A0R612, A0R613, A0R614, A0R616, A0R617, A0R618, A0R619, A0R620, A0R623, A0R624, A0R625, A0R626, A0R627, A0R628, A0R629, A0R632, A0R633, A0R634, A0R636, A0R637, A0R638, A0R639, A0R640, A0R642, A0R643, A0R646, A0R647, A0R648, A0R649, A0R650, A0R652, A0R655, A0R656, A0R659, A0R663, A0R664, A0R665, A0R666, A0R670, A0R676, A0R678, A0R679, A0R683, A0R684, A0R689, A0R692, A0R696, A0R698, A0R699, A0R6A0, A0R6A2, A0R6A3, A0R6A4, A0R6A5, A0R6A6, A0R6A7, A0R6A8, A0R6A9, A0R6B0, A0R6B1, A0R6C4, A0R6C7, A0R6C8, A0R6D1, A0R6D2, A0R6D3, A0R6D4, A0R6D6, A0R6D7, A0R6D9, A0R6E0, A0R6E3, A0R6E4, A0R6E5, A0R6E6, A0R6E9, A0R6F1, A0R6G3, A0R6G6, A0R6G7, A0R6H1, A0R6H7, A0R6H8, A0R6I1, A0R6I2, A0R6I4, A0R6I9, A0R6J8, A0R6J9, A0R6K1, A0R6K4, A0R6K6, A0R6K8, A0R6K9, A0R6L2, A0R6L4, A0R6L5, A0R6L6, A0R6L7, A0R6L8, A0R6L9, A0R6M0, A0R6M1, A0R6M2, A0R6M4, A0R6M5, A0R6M6, A0R6M7, A0R6N5, A0R6N9, A0R6P0, A0R6P8, A0R6P9, A0R6Q0, A0R6Q3, A0R6Q7, A0R6Q9, A0R6R0, A0R6R4, A0R6R5, A0R6R6, A0R6R9, A0R6S0, A0R6T1, A0R6T2, A0R6T3, A0R6T5, A0R6U3, A0R6V5, A0R6V6, A0R6V7, A0R6W0, A0R6W5, A0R6Z0, A0R6Z3, A0R6Z4, A0R6Z5, A0R6Z6, A0R6Z9, A0R703, A0R708, A0R710, A0R711, A0R712, A0R713, A0R716, A0R717, A0R718, A0R720, A0R722, A0R723, A0R724, A0R726, A0R727, A0R729, A0R730, A0R731, A0R742, A0R746, A0R748, A0R749, A0R750, A0R753, A0R757, A0R758, A0R759, A0R763, A0R768, A0R771, A0R772, A0R773, A0R775, A0R776, A0R782, A0R788, A0R795, A0R7A8, A0R7C5, A0R7C8, A0R7C9, A0R7D0, A0R7D8, A0R7D9, A0R7E1, A0R7E4, A0R7E5, A0R7E9, A0R7F4, A0R7F6, A0R7F7, A0R7F9, A0R7G2, A0R7G4, A0R7G5, A0R7G6, A0R7G7, A0R7G8, A0R7H0, A0R7H1, A0R7H2, A0R7H3, A0R7H4, A0R7H5, A0R7H6, A0R7H7, A0R7H8, A0R7H9, A0R7I0, A0R7I2, A0R7I3, A0R7I4, A0R7I5, A0R7I9, A0R7J0, A0R7J1, A0R7J2, A0R7J3, A0R7J4, A0R7J5, A0R7J6, A0R7J7, A0R7K1, A4ZHR8, A4ZHT6, O08323, O52199, O52200, O68956, O85501, P0CH00, P0CG99, P0CH37, P0CH36, P42829, P48354, P60281, P71533, P71534, P94968, Q2M5K2, Q2M5K3, Q2M5K4, Q2YHI9, Q3I5Q7, Q3L885, Q3L887, Q3L890, Q3L891, Q50441, Q59560, Q9AFI5, Q9F868, Q9RMN9, Q9RP48, Q9X5M0, Q9X5M1, Q9ZHC5, A0R091, A0QYS5, A0QTX2, A0QY48, A0QUZ6, A0QR99, A0QZ46, A0R635, A0QQW4, A0R0C7
