## Supplementary File S2 - Preselected dataset for "OrthoGather: a local platform for orthology-based proteome comparisons and Gene Ontology enrichment"

**Supplementary File S2. List of reference proteomes available through the Preselected Dataset input mode.** Forty-seven bacterial proteomes from 35 species across 19 genera, bundled with OrthoGather and initially assembled for studies of cystic fibrosis associated and highly antibiotic-resistant pathogens. **No:** numerical index of the entry; **Strain:** organism and strain designation as given in UniProt; **NCBI ID:** NCBI Taxonomy identifier; **Proteome ID:** UniProt proteome accession.

| No | Strain | NCBI ID | Proteome ID |
|---|---|---|---|
| 1 | *Achromobacter insuavis* AXX-A | 1003200 | UP000004853 |
| 2 | *Achromobacter ruhlandii* LMG 3328 | 72557 | UP000494122 |
| 3 | *Achromobacter xylosoxidans* C54 | 562971 | UP000006414 |
| 4 | *Acinetobacter baumannii* AB5075 | 1116234 | UP001220259 |
| 5 | *Acinetobacter baumannii* AB900 | 557601 | UP000762945 |
| 6 | *Acinetobacter baumannii* ATCC 17978 | 400667 | UP000072389 |
| 7 | *Acinetobacter baumannii* ATCC 19606 | 575584 | UP000005740 |
| 8 | *Acinetobacter baumannii* AYE | 509173 | UP000002446 |
| 9 | *Acinetobacter baylyi* ADP1 | 62977 | UP000000430 |
| 10 | *Acinetobacter calcoaceticus* P23 | 471 | UP000031220 |
| 11 | *Burkholderia cenocepacia* J2315 | 216591 | UP000001035 |
| 12 | *Burkholderia dolosa* FDAARGOS_1272 | 152500 | UP000625568 |
| 13 | *Burkholderia gladioli* BSR3 | 999541 | UP000008316 |
| 14 | *Burkholderia multivorans* ATCC 17616 | 395019 | UP000008815 |
| 15 | *Burkholderia pseudomallei* K96243 | 272560 | UP000000605 |
| 16 | *Burkholderia stabilis* FERMP-21014 | 95485 | UP000218432 |
| 17 | *Burkholderia thailandensis* E264 | 271848 | UP000263968 |
| 18 | *Burkholderia vietnamiensis* G4 (LMG 22486) | 269482 | UP000002287 |
| 19 | *Chryseobacterium indologenes* CI_885 | 253 | UP000037953 |
| 20 | *Cupriavidus respiraculi* LMG 21510 | 195930 | UP000721236 |
| 21 | *Enterococcus faecium* | 1352 | UP000873179 |
| 22 | *Escherichia coli* K-12 MG1655 | 83333 | UP000000625 |
| 23 | *Escherichia coli O157:H7 str. Sakai* | 83334 | UP000000558 |
| 24 | *Haemophilus influenzae* Rd KW20 (ATCC 51907) | 71421 | UP000000579 |
| 25 | *Inquilinus limosus* Inq | 171674 | UP000196655 |
| 26 | *Klebsiella oxytoca* | 571 | UP000873349 |
| 27 | *Klebsiella pneumoniae* subsp. *pneumoniae* HS11286 | 1125630 | UP000007841 |
| 28 | *Mycobacterium avium* subsp. *hominissuis* MAC109 | 439334 | UP000259236 |
| 29 | *Mycobacterium intracellulare* ATCC 13950 | 487521 | UP000008004 |
| 30 | *Mycobacterium tuberculosis* H37Rv | 83332 | UP000001584 |
| 31 | *Mycobacteroides abscessus* ATCC 19977 | 561007 | UP000007137 |
| 32 | *Mycobacteroides abscessus* subsp. *bolletii* BD | 1091046 | UP000005975 |
| 33 | *Mycobacteroides abscessus* subsp. *massiliense* 787 | 1962118 | UP000189730 |
| 34 | *Mycobacteroides chelonae* CCUG 47445 | 1460372 | UP000076804 |
| 35 | *Mycolicibacterium smegmatis* mc$^2$155 (ATCC 700084) | 246196 | UP000000757 |
| 36 | *Nocardia farcinica* IFM 10152 | 247156 | UP000006820 |
| 37 | *Pandoraea sputorum* NCTC 13161 | 93222 | UP000215126 |
| 38 | *Pseudomonas aeruginosa* 1BAE | 287 | UP001190961 |
| 39 | *Pseudomonas aeruginosa* LESB58 | 557722 | UP000001527 |

| No | Strain | NCBI ID | Proteome ID |
|---|---|---|---|
| 40 | *Pseudomonas aeruginosa* PA103 | 1081927 | UP001302188 |
| 41 | *Pseudomonas aeruginosa* PA14 | 652611 | UP001061545 |
| 42 | *Pseudomonas aeruginosa* PAK | 1009714 | UP000014100 |
| 43 | *Pseudomonas aeruginosa* PAO1 | 208964 | UP000002438 |
| 44 | *Pseudomonas putida* KT2440 | 160488 | UP000000556 |
| 45 | *Ralstonia mannitolilytica* R-77591 | 105219 | UP001190002 |
| 46 | *Staphylococcus aureus* NCTC 8325 (PS 47) | 93061 | UP000008816 |
| 47 | *Stenotrophomonas maltophilia* K279a | 522373 | UP000008840 |
